# Co-aggregation and secondary nucleation in the life cycle of human prolactin/galanin functional amyloids

**DOI:** 10.1101/2021.08.31.458467

**Authors:** D. Chatterjee, R.S. Jacob, S. Ray, A. Navalkar, N. Singh, S. Sengupta, L. Gadhe, P. Kadu, D. Datta, A. Paul, C. Pindi, S. Kumar, P. S. Singru, S. Senapati, S. K. Maji

## Abstract

Synergistic-aggregation and cross-seeding by two different amyloid proteins/peptides are well evident in various neurological disorders. However, this phenomenon is not well studied in functional amyloid aggregation. Here, we show Prolactin (PRL) is associated with lactation in mammals and neuropeptide galanin (GAL), which are co-stored in the lactotrophs facilitates the synergic aggregation in the absence of secretory granules helper molecules glycosaminoglycans (GAGS). Interestingly, although each partner possesses homotypic seeding ability, a unidirectional cross-seeding of GAL aggregation can be mediated by PRL seeds. The specificity of co-aggregation by PRL and GAL along with unidirectional cross-seeding suggests tight regulation of functional amyloid formation during co-storage of these hormones in secretory granule biogenesis of female rat lactotrophs. Further mixed fibrils release the constituent functional hormone much faster than the corresponding individual amyloid formed in presence of GAGs, suggesting that co-aggregation of functionally distant hormones might have evolved for efficient storage, synergistic and rapid release of both hormones upon stimulation. The co-aggregation and cross seeding by two different hormones of completely different structures and sequences (PRL and GAL) suggest a novel mechanism of heterologous amyloid formation both in disease and functional amyloids.

## Introduction

Protein/peptide misfolding, aggregation, and amyloid formation is associated with various neurological disorders such as Alzheimer’s and Parkinson’s^1,2^. However, several studies have suggested that amyloid formation is also associated with the native biological function of the host organism. The protein fibrils formed to aid in the functionality of the host organisms are called functional amyloids^3-7^. For example, Pmel-17 amyloid fibrils template melanin polymerization inside melanosomes^8-10^; Curli fibrils of *E. coli* support the organism to adhere to a surface and also help their colonization process inside biofilms^11,12^. In this line, another very interesting aspect is the formation of functional amyloid by protein/peptide hormones (such as prolactin (PRL) and galanin (GAL)) during their storage inside the secretory granules (SGs) of pituitary^13,14^. The amyloid formation, in this case, not only enriches the protein concentration to serve as a protease-resistant protein/peptide storage depot but is also able to release the functional monomeric proteins upon dilution and pH changes^13-17^.

Protein/peptide aggregation and amyloid formation generally follow nucleation-dependent polymerization mechanism^18-20^, where protein/peptide slowly associates to form aggregation competent nuclei (in the lag phase of aggregation)^21-23^. Once formed, the aggregation competent nuclei further recruit the monomeric counterpart for their growth into mature amyloid fibrils (elongation phase)^21-23^. The progression of aggregation eventually reaches a steady-state equilibrium between the fibrils and monomeric protein (stationary phase)^19,24^. Recent evidences however suggest that fragmentation/elongation and secondary nucleation may contribute to a significant decrease in the lag-time of aggregation, similar to an external addition of preformed nuclei in amyloid growth reaction (seeding)^25-27^. Although homotypic aggregation and seeding is the most favored mechanism of protein aggregation and amyloid formation, synergistic aggregation (co-aggregation) by two different proteins/peptides and heterologous seeding are also suggested to be involved in many neurodegenerative disorders^28-32^. This is one of the possible mechanisms by which one disease aggravates the other disease such as Alzheimer’s disease and Type 2 diabetes and Alzheimer’s disease and Parkinson’s disease^33,34^.

PRL and GAL secretion is synergistic and promoted by common secretagogues^35-37^. PRL/GAL co-storage has been also reported in the anterior pituitary of female rats or estrogen-treated male rats^38^. Here, we investigate the synergistic aggregation and amyloid formation by these hormones for their secretory storage and release from secretory granules. Indeed, both of these hormones are co-stored in the female rat pituitary and possess amyloid-like characteristics. In this study, we show that both hormones not only engage in homotypic amyloid aggregation in the presence of respective GAGs as helper molecules but also synergistically aggregate (in absence of GAGs) to form heterotypic amyloid containing PRL and GAL as suggested by double immunoelectron microscopy. Intriguingly, cross-seeding with PRL fibril seeds resulted in fibrillation of GAL. However, GAL fibrils could not seed PRL monomers for fibrillation. This suggests a tightly controlled regulation of hormone amyloids for their homotopic and heterotypic storage. Furthermore, our *in vitro* release assay showed faster release of functional monomers by heterotypic, hybrid amyloid (PRL+GAL) compared to homotypic counterparts. This supports the storage and release are highly controlled and conserved in pituitary tissue for the optimum function to be served.

## Results

### PRL and GAL are co-stored as amyloids in SGs of the anterior pituitary of female rat

Previously it was shown that PRL and GAL co-store in the anterior pituitary of female rats or estrogen-treated male rats^38-40^. PRL is a 23 kDa protein (**Supplementary Table 1**) and consists of four-helix bundles comprising residues 14-42 (helix 1), 78-104 (helix 2), 110-138 (helix 3) and 160-194 (helix 4)^41,42^. The helix bundles are spaced with 3 loop regions (**Fig 1a**). On the other hand, GAL is a small unstructured neuropeptide (**Supplementary Table 1**), which is 30 residues in length^43,44^ (**Fig 1a**). PRL showed amyloid-prone sequences predicted by TANGO^45^; GAL, however, did not contain any amyloid-prone sequences (**Fig 1b-c**). Since previously it was established that protein/peptide hormones (including PRL and GAL) can be stored as amyloids inside the (SGs)^13^, we tested whether PRL and GAL are colocalized at anterior pituitary in the amyloid state or not. To examine this possibility, immunofluorescence study of female rat pituitary was performed using anti-GAL (green) and anti-PRL (red) antibodies (see ***material and methods***). Our results showed that PRL and GAL were substantially co-localized in the pituitary tissue (**Fig 1d**) indicating their co-storage inside the SGs. Further, to test whether both PRL and GAL remain in the amyloid state, we performed immunostaining of the tissue with amyloid specific antibody OC along with Thioflavin-S (ThioS)^46^ (**Fig 1e, Supplementary Figure 1**). Double immunofluorescence of PRL and GAL along with amyloid specific OC antibody showed strong colocalization suggesting both the hormones are in the amyloid state (**Fig 1e**), which was also observed in ThioS staining (**Supplementary Figure 1**). Altogether, our data confirmed that both PRL and GAL are co-stored as amyloid aggregates in the female rat pituitary.

**Fig 1:**
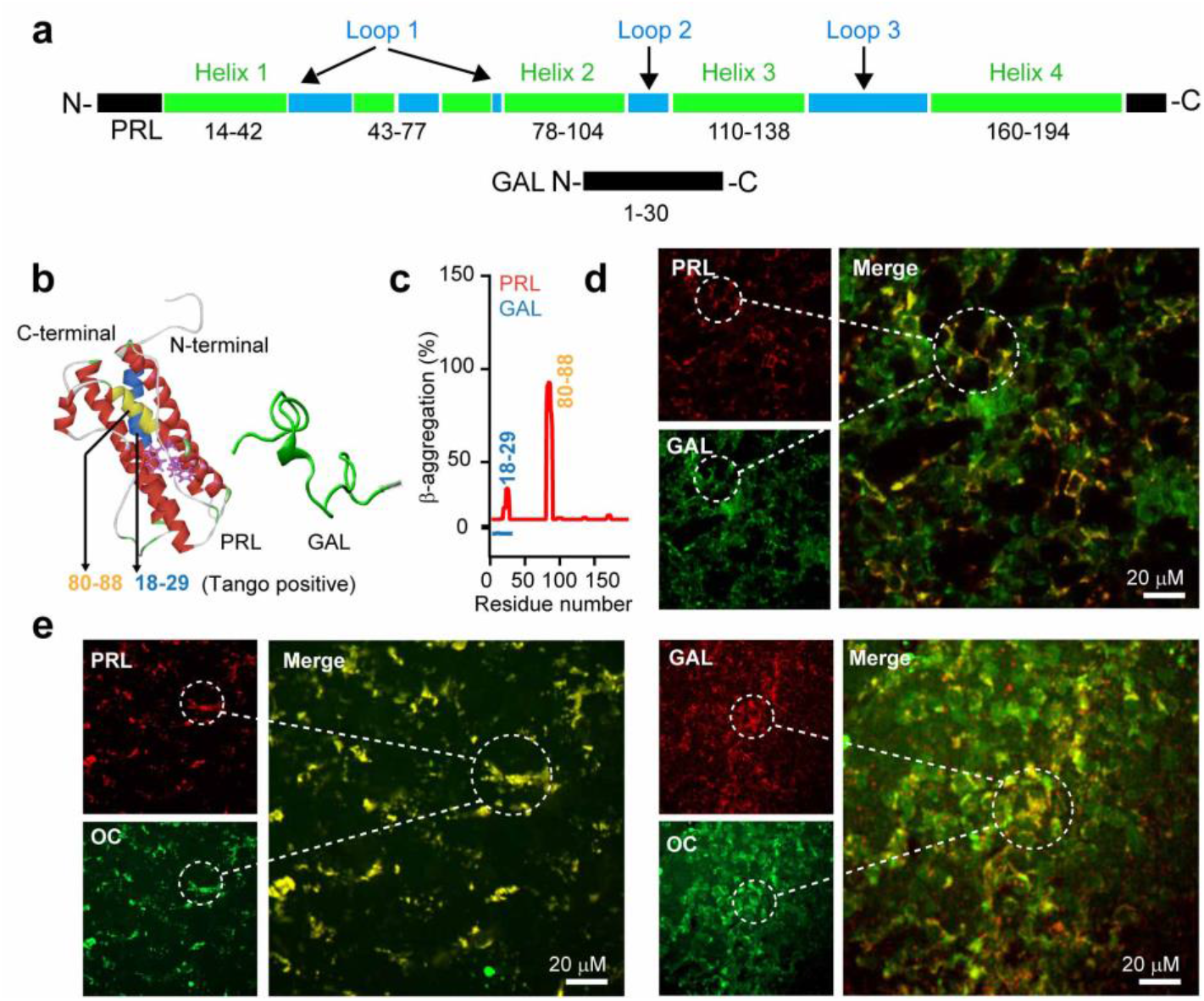
Amyloid propensity and co-storage of PRL and GAL. **a**. Schematic showing amino acid sequence and secondary structures of PRL & GAL with different color codes. (***Upper panel***): PRL is 191 amino acids in length and contains a four-helix bundle (green). The short helix and loop regions are also represented between helix-1 and helix 2 (shown in green and blue colors, respectively). (***Lower panel***): GAL showing 30 residue peptide with no definite secondary structure^43,44^. **b. (*Left panel*)** The three-dimensional structure (obtained in Pymol)^41^ of PRL showing its major helices and two tryptophan residues (shown in purple) (***PDB ID: 1RW5***). **(*Right panel*):** Natively unstructured conformation of GAL is also shown. **c**. TANGO algorithm showing the aggregation-prone residues of PRL and GAL at pH 6.0 (SG relevant pH). The residues 18-29 and 80-88 of PRL showing amyloid aggregation potential. However, TANGO analysis of GAL revealed no amyloid aggregation propensity. Immunofluorescence studies showing (**d**) colocalization of PRL (red) and GAL (green) in the female rat anterior pituitary. (**e) *left panel***. Colocalization of amyloid fibrils (OC, green) and PRL (red) and amyloid fibrils (OC, green) and GAL (red) (***Right panel***)) in the anterior pituitary. The merged photomicrographs showing colocalization (yellow). The experiments (**d-e**) are performed three times with similar observations.

### *In vitro* amyloid aggregation kinetics and synergistic co-fibrils of PRL and GAL

The colocalization of both PRL and GAL in the amyloid state suggested that PRL and GAL might interact with each other and might co-aggregate inside the same SGs. We hypothesized that there could be 4 possibilities (case 1-4) when PRL and GAL are co-stored as amyloids (**Fig 2a**). The two proteins can form separate filaments and these filaments can be incorporated into heterogeneous fibrils (case 1). They can form completely separate, homogeneous fibrils either by PRL or GAL or by both hormones (case 2). It is also possible that the PRL and GAL together can form the fibril forming unit to form heterogeneous amyloid fibrils (case 3); and lastly, the PRL monomers can simply adhere to the GAL fibril (case 4) and vice-versa (case 5).

**Fig 2:**
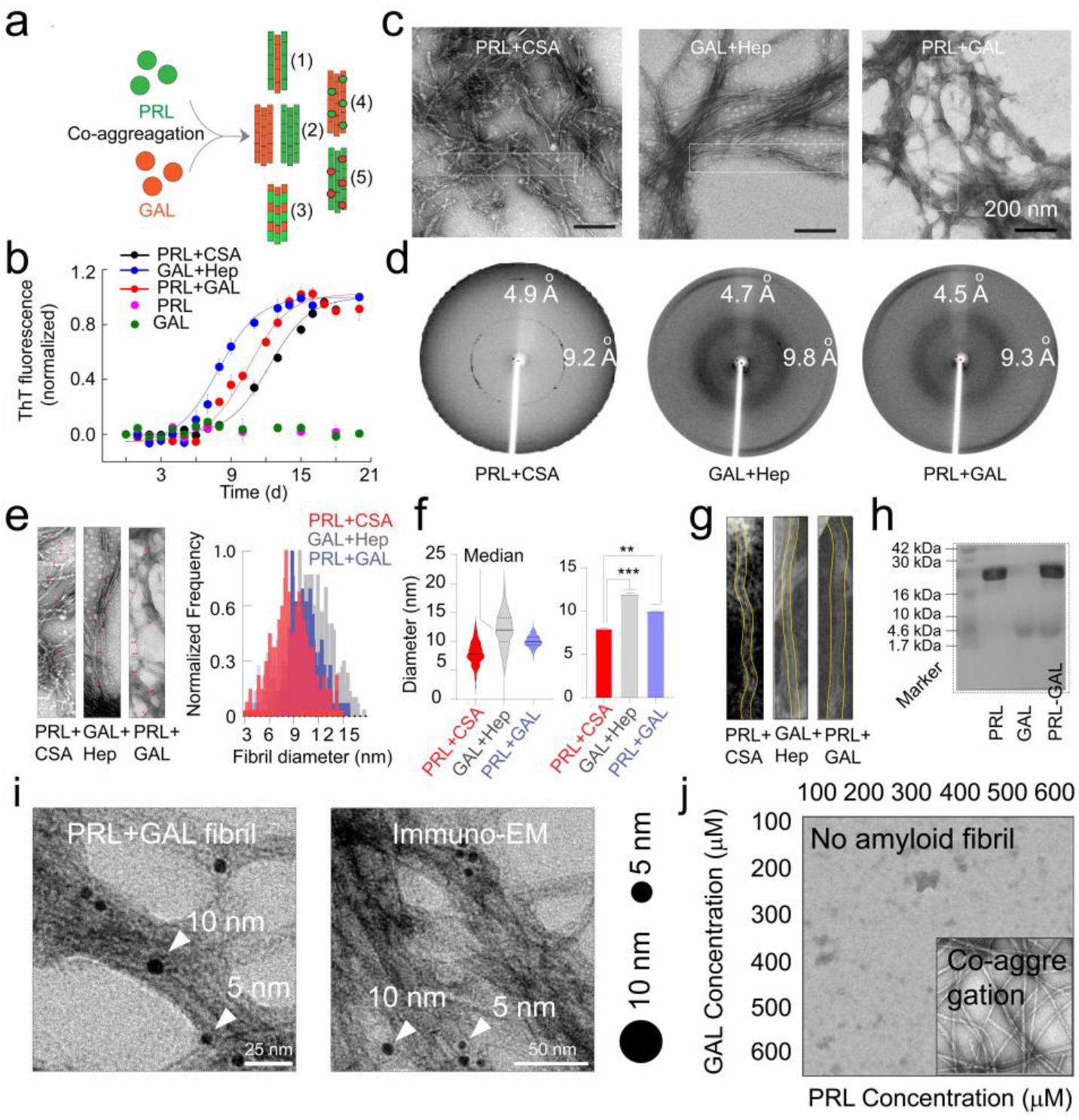
Amyloid aggregation kinetics & co-fibril formation by PRL and GAL: **a**. Schematic showing different possibilities for the formation of PRL-GAL co-fibril. **b**. Normalized ThT fluorescence intensity over time showing faster aggregation kinetics for GAL-Hep followed by PRL-GAL and PRL-CSA. The experiment is performed three times with similar results. Values represent mean ± SEM. **c**. TEM images showing amyloid fibrils for PRL-CSA, GAL-Hep, and PRL-GAL after 15 days of incubation. Representative images are shown. The dotted box marks are the representative area from which the fibril diameters are measured. **d**. XRD of PRL-CSA, GAL-Hep, and PRL-GAL fibrils at day 15 showing ∼4.7 Å meridional and ∼10 Å equatorial reflections, as commonly seen for most amyloid fibrils^47,48^. **e**. (***Left panel***) Representative TEM images showing fibril diameter measured at random positions (marked with red arrows) on individual fibrils. (***Right panel***) The normalized frequency distribution of fibril diameters of PRL+CSA, GAL+Hep, and PRL+GAL fibrils is shown. 200 data points are collected for individual samples for n=3 independent experiments. **f. (*Left panel*)** Median values of different fibril diameters are shown with violin plots. (***Right panel***) Average values of different fibril diameters are shown. Values represent mean ± SD for n=3 independent experiments. The statistical significance (***p ≤ 0.001, **p ≤ 0.01) is calculated by one-way ANOVA followed by an SNK post hoc test with a 95% confidence interval. **g**. Representative TEM images of PRL-CSA, GAL-Hep, and PRL-GAL fibrils (scale bar-200 nm). From a single fibril, 200 data points are collected along the length to calculate the diameter. **h**. SDS-PAGE depicting two bands for isolated aggregates from the PRL-GAL mixture (lane 3). The two bands correspond to PRL and GAL, which suggests that the isolated aggregates are composed of both PRL and GAL. **i**. Amyloid fibrils obtained from the PRL-GAL mixture showing 10 nm gold particles (against GAL primary) and 5 nm gold particles (against PRL primary), confirming synergistic co-fibril formation by PRL and GAL (***Left and Right panel***). The experiment is performed three times with similar observations. **j**. Schematic representation of incubation of PRL and GAL at various concentrations showing optimum concentration is required to initiate PRL-GAL co-aggregation.

To explore these possibilities, we asked whether two hormones with completely different sequences, length and without any sequence similarity/identity (**Supplementary Figure 2**) can co-assemble to form hybrid fibrils. To do this, we mixed the 1:1 molar ratio of both the hormones, incubated at 37 °C, and followed the amyloid formation by Thioflavin-T (ThT) binding and CD spectroscopy for 15 days (**Fig 2b, Supplementary Figure 2**). As a control, both hormones were incubated alone. Since PRL and GAL are known to form amyloid in presence of chondroitin sulfate A (CSA) and Heparin (Hep)^13^, we also incubated both the hormones in presence of their respective GAGs as positive controls (**Fig 2b, Supplementary Figure 2**). As expected, both PRL and GAL showed fibril formation in the presence of CSA and Hep. No ThT binding was observed by either PRL or GAL in absence of GAGs (**Fig 2b, Supplementary Figure 2**).

Strikingly, the mixture of both the hormones showed high ThT binding during time indicating amyloid formation (**Fig 2b, Supplementary Figure 2**).CD spectroscopy showed a substantial decrease of helical content by PRL-GAL and PRL-CSA during aggregation. Notably, PRL is known to form amyloid without structural conversion to β-sheet^13^. On the other hand, GAL-Hep showed random coil to β-sheet conversion during amyloid formation (**Supplementary Figure 2**). The lag times of aggregation for PRL-CSA, PRL-GAL, and GAL-Hep were calculated as ∼9.2 days, ∼8.8 days, and ∼7 days, respectively (**Supplementary Figure 2**), suggesting that PRL-GAL aggregated faster than PRL-CSA. Moreover, PRL-GAL aggregates showed fibril like morphology under TEM (**Fig 2c**), strong apple-green/golden birefringence under cross-polarized light (**Supplementary Figure 3**), exhibited cross β-sheet diffraction patterns^47,48^ (∼4.7Å for inter-strand and ∼9.8 Å for inter-sheet) (**Fig 2d**) and FTIR peaks corresponding to β-sheet structure^49,50^ (**Supplementary Figure 4**). Similar observations supporting amyloid structure were also obtained for PRL-CSA and GAL-Hep aggregates (**Fig 2c-d** and **Supplementary Figures 2-4**).

Although there are various possibilities regarding how PRL-GAL can form fibrils (any of the component hormones can form fibrils separately or together, as described in **Fig 2a**), we analyzed the frequency distribution of the fibril diameters from TEM images. Fibril diameters were measured from a particular sample (n=200 randomized points) (**Fig 2e, *left panel***) and the normalized frequency distribution was plotted (**Fig 2e, *right panel***). Our data indicated that the diameter of PRL-CSA fibrils was least (**Fig 2e, *right panel***) having a median value of ∼7.5 nm (**Fig 2f, *left panel***) and an average of ∼7.8 nm (**Fig 2f, *right panel***). The diameter of GAL-Hep fibrils was highest (**Fig 2e, *right panel***) having a median value of ∼12.3 nm (**Fig 2f, *left panel***) and an average of ∼12 nm (**Fig 2f, *right panel***). Intriguingly, the PRL-GAL fibrils showed an intermediate diameter (median∼10 nm, average∼9.8 nm) (**Fig 2e-f**). This data suggest that PRL-GAL fibrils might be a new type of hybrid fibrils, which is neither similar to GAL nor PRL fibrils. We further analyzed the diameter from the same fibril bundle along its length. Consistent with our random point analysis (**Fig 2e-f**), we found that the diameter of a single PRL-CSA fibril was indeed the lowest (average∼7 nm) followed by PRL-GAL (average∼10 nm) and GAL-Hep (average∼12 nm) fibril (**Fig 2g, Supplementary Figure 2**). Our data indicate that PRL-GAL fibrils possess unique morphology, which could be due to the incorporation of both PRL as well as GAL molecules into the same fibril (co-fibril).

To further analyze whether both PRL and GAL are part of the same insoluble fibril fraction, we isolated the fibrils using centrifugation and performed SDS-PAGE. The presence of two bands corresponding to PRL and GAL indicated that both PRL and GAL formed fibrils when co-incubated together (**Fig 2h**). Further, we performed immuno-electron microscopy (Immuno EM) with the PRL-GAL fibrils in the presence of secondary antibodies against PRL and GAL attached with 5 nm and 10 nm gold nanoparticles, respectively (**Fig 2i**). Co-localization of both 5 and 10 nm nanoparticles within the same fibril bundle confirmed that both PRL and GAL were part of the same fibril bundle (**Fig 2i**). However, due to extensive bundling of fibrils, it remains to be seen whether PRL and GAL are incorporated in the same filament or not.

Since 400 µM (each) PRL-GAL could undergo aggregation even in the absence of any helper molecules, we wanted to understand the optimum concentration and stoichiometry for PRL-GAL co-aggregation. To do this, we chose increasing concentration of PRL and GAL in an orthogonal (X-Y axis) manner (from 100-600 µM, each) (**Fig 2j, Supplementary Fig 5**). All combinations of this concentration regime of hormone mixture were incubated for 15 days. We found≥1:1 ratio and ≥400 µM concentration are required for mixed fibrils formation (experimentally feasible time scale) as determined by ThT binding and electron microscope study (**Fig 2j, Supplementary Fig 5**). The data suggest that the equimolar ratio of PRL and GAL can co-assemble to form hybrid fibrils in a concentration-dependent manner.

### Cross-seeding of PRL and GAL

Amyloid aggregation is a nucleation-dependent polymerization process^18,19^. It is well-known fact that the presence of preformed nuclei of amyloid (also called “seeds”) greatly affects the kinetics of aggregation of monomers^25,51^. Since PRL and GAL co-aggregate into mixed amyloids, we asked whether both PRL and GAL could cross-seed^25,51-54^ to induce amyloid aggregation of each other and help their possible storage in SGs. To do that, we performed both homotypic (PRL monomer+PRL seed; GAL monomer+GALseed) and heterotypic seeding (PRL monomer+GAL seed and GAL monomer+PRL seed).

Preformed fibrils were sonicated to obtain PRL and GAL amyloid fibril seeds (see ***materials and methods***). 1%, 2%, and 5%(v/v) of PRL and GAL seeds were mixed with freshly prepared 400 µM PRL and GAL, respectively, and incubated with slight agitation at 37°C for homotypic seeding (**Fig 3b, Supplementary Figure 6**). Our ThT fluorescence data showed accelerated aggregation by both PRL and GAL for homotypic seeding as lag time decreased significantly in the presence of 2% and 5% seeds (**Fig 3b, 3e**). However, 1% seed did not show any fibril formation for both PRL as well as GAL (**Fig 3b, Supplementary Figure 6**) even after 10 days of incubation. Fibril formation via homotypic seeding mechanism was further supported by TEM imaging and time-dependent CD spectroscopic measurements (**Fig 3b-c, Supplementary Figure 6**).

**Fig 3:**
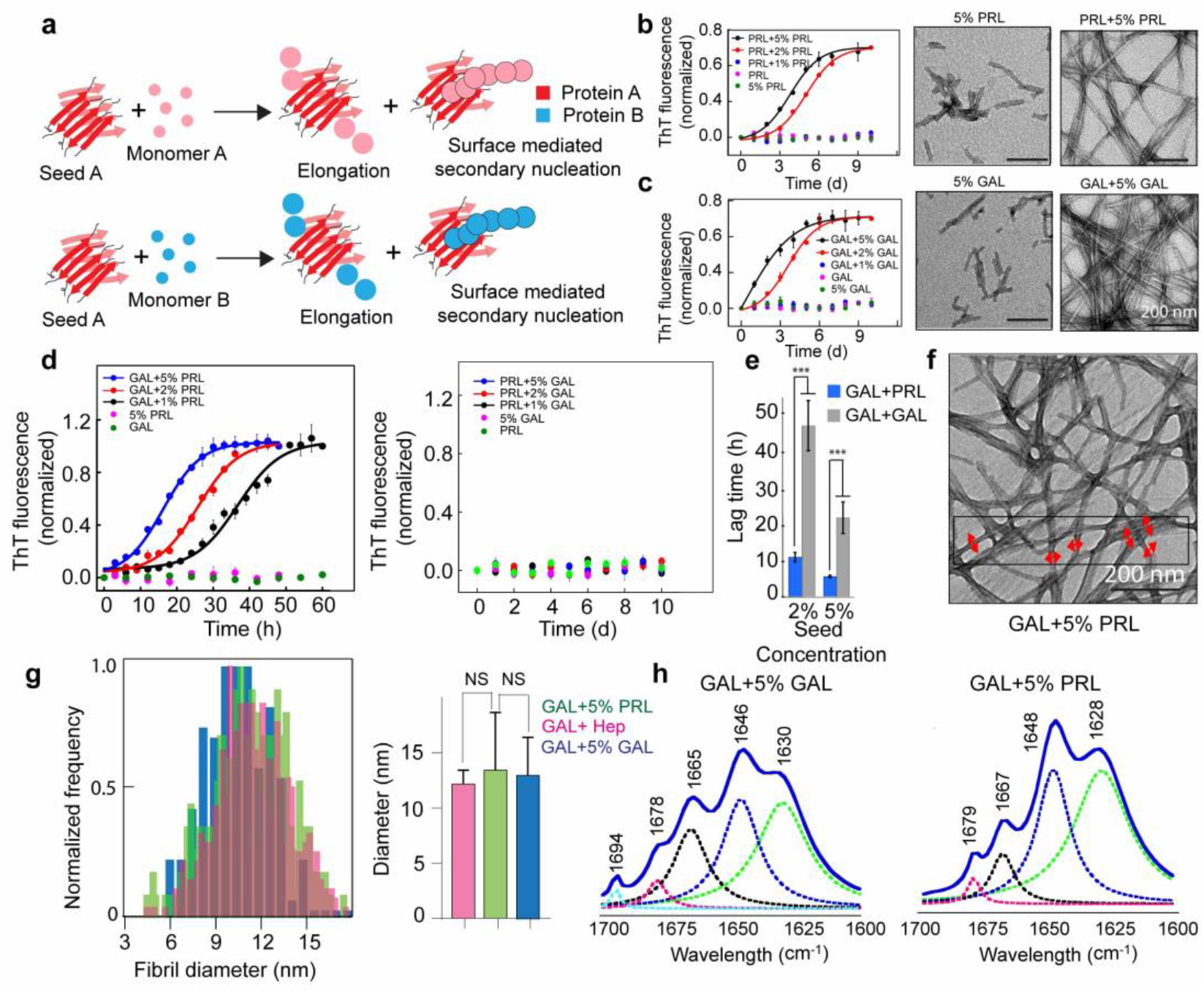
Seeding and cross-seeding of PRL and GAL: **a**. Schematic showing possible homo and hetero seeding with fibril elongation and surface-mediated secondary nucleation mechanism for seed-mediated fibril growth. **(b-c). homo-seeding of PRL and GAL**. (***left panel***) PRL and GAL homo-seeding. Normalized ThT fluorescence intensity values with time indicating aggregation of PRL and in the presence of different concentrations of PRL seeds and GAL seeds respectively (2% and 5% v/v). Only seeds and only PRL/GAL was used as controls. (***right panel***) The corresponding EM images of PRL/GAL seeds alone and PRL/GAL monomer in presence of 5% PRL/GAL seeds showing fibrils formation by PRL/GAL homo-seeding. **d. Cross-seeding of PRL and GAL**. (***left panel*)** Normalized ThT fluorescence intensity values with time indicating aggregation of GAL in the presence of different concentrations of PRL seeds (1%, 2%, and 5% v/v). However, PRL in presence of different percentages of GAL seeds does not show any aggregation (***right panel***). Only seed and only GAL/PRL were used as controls where no aggregation is observed. e. The lag times of GAL aggregation in presence of 2% and 5% v/v PRL seeds and GAL seeds are compared. The values represent mean ± SEM. The significance (***p ≤ 0.001) is calculated using one-way ANOVA followed by an SNK post hoc test with a 95% confidence interval. **f**. TEM images of GAL fibrils formed in presence of PRL seeds are shown. GAL fibrils formed in presence of 5% (v/v) PRL seeds are analyzed for frequency distribution (red arrows indicating the diameter of the fibrils measured for analysis). **g. (*left panel*)** Normalized frequency distribution of fibril diameter showing GAL fibrils formed in presence of PRL seeds have a similar diameter to GAL+Hep fibrils. 200 random data points from different individual fibrils were collected from n=3 independent experiments for the frequency distribution analysis. (***right panel***) Average values of different fibril diameters are shown. Values represent mean ± SD for n=3 independent experiments. The statistical significance (***p ≤ 0.001, **p ≤ 0.01) is calculated by one-way ANOVA followed by an SNK post hoc test with a 95% confidence interval. **h**. FTIR spectra showing fibrils of GAL+5% GAL seed and GAL+5% PRL seed are of similar secondary structure.

Similar experiments were done where various concentrations of PRL seeds were mixed with GAL monomer and GAL seeds were mixed with PRL monomer (heterotypic seeding). We hypothesized two possibilities of secondary nucleation^55^—the heterotypic monomers can be recruited at the ends of the seeds and grow the amyloid fibril (elongation) or the seed surface will help in the nucleation of the heterotypic monomer^55^ (**Fig 3a**). Our data showed that GAL aggregation was accelerated with PRL seeds in a concentration-dependent manner (**Fig 3d, *left panel***). Surprisingly, we observed no ThT fluorescence for all seed concentrations even after 10 days of incubation when GAL seeds were incubated with PRL monomers, indicating that GAL fibrils are incapable of inducing amyloid fibril formation of PRL monomers (**Fig 3d, *right panel***). This was also evident with CD spectroscopic measurements (**Supplementary Fig 7**). Interestingly, no aggregation was observed when PRL/GAL monomer was incubated in presence of different percentages of PRL-GAL mixed fibril seeds (1%, 2%, and 5%) as confirmed by CD spectroscopy, ThT fluorescence, and TEM imaging (**Supplementary Fig 8-9**). This suggests that seeding (both homo and hetero) event is very specific for the life cycle of PRL/GAL amyloid formation in SG. Important to note that these homotypic and heterotypic seeding studies were done in the absence of any helper molecules of GAGs. When lag times were compared, the heterotypic seeding rate was significantly higher than homotypic seeding for GAL aggregation suggesting surface-mediated secondary nucleation might be triggering GAL aggregation in the presence of PRL seeds^56^ (**Fig 3e**).

To further understand whether PRL seeds engage the GAL monomer for secondary nucleation^27^ where elongation and/or surface-mediated aggregation could happen^55^, fibril diameters of GAL (formed by PRL cross-seeding) were analyzed from the TEM images (**Fig 3f**). If the elongation/fragmentation mechanism is occurring, GAL will form PRL-like fibrils. If PRL seeds engage GAL monomer for surface-mediated secondary nucleation, GAL will essentially form GAL-like fibrils. Intriguingly, analysis of frequency distribution of the GAL fibril diameters indicated that the distribution of diameter of PRL seeded GAL fibrils was very similar to that of GAL-Hep fibrils (**Fig 3g, *left panel***). The median and mean fibril diameter analysis also suggests that GAL fibril formed in presence of PRL seed (median and mean diameter 12.35 nm) is very similar compared to GAL fibrils formed in presence of GAL seed (median and mean diameter 12 nm) as well as GAL fibrils formed in presence of heparin (median and mean diameter 11.20 nm) (**Fig 3g, *right panel*, Supplementary Figure 6**). This means that the incorporation of GAL monomers does not happen to the PRL fibril end (elongation)^25,51,57,58^, rather, GAL monomers use PRL seeds as surface and form amyloid fibrils via secondary (or surface) nucleation^26,52,53,56^ mechanism. This was further confirmed with the FTIR study, which suggested that GAL-Hep and GAL in the presence of PRL seed possessed identical FTIR spectral signature, which was substantially different from the PRL fibril spectrum (**Fig 3h**).

### Specific interactions of PRL and GAL leading to amyloid aggregation

Next, we wanted to further investigate if interactions leading to co-aggregation and amyloid formation of PRL and GAL are specific to themselves. To do this, we co-incubated PRL with adrenocorticotropic hormone (ACTH) (**Supplementary Table 1**) as this hormone is of similar length to GAL and does not form amyloid by itself ^13,59^. Similarly, GAL was also incubated with growth hormone (GH) (**Supplementary Table 1**), a hormone structurally and functionally related to PRL^60,61^. ThT aggregation kinetics and CD spectroscopy were performed at the beginning of the aggregation (day 0) and after 15 days of incubation for both PRL-ACTH and GAL-GH. Our data showed negligible ThT fluorescence for both PRL-ACTH and GAL-GH even after 15 days (**Fig 4a**) suggesting no co-aggregation. This observation was consistent with no structural conversion observed in CD and TEM, where the PRL-ACTH and GH-GAL mixtures were devoid of any fibrils (**Fig 4b, Supplementary Figure 10**). Overall, our observations suggest that interaction and co-aggregation/amyloid formation by PRL and GAL are specific and are mutually beneficial for the storage of these hormones in SGs. This specific co-aggregation of PRL and GAL could be due to their favorable interaction when the monomeric hormone is mixed together. This is further evident from the surface plasmon resonance (SPR) study, where GAL monomers were immobilized on a CM-5 chip and a range of concentrations of PRL monomeric protein was passed over it. We observed a significant increase in the response unit (RU) indicating binding of PRL to GAL (**Fig 4c**). The relative dissociation constant (K_D_) was calculated to be 4.1×10^−7^ M, which indicates the binding of PRL with GAL (**Fig 4d**). In comparison, we observed no significant binding when ACTH was passed through immobilized PRL or when GH was passed through immobilized GAL (**Fig 4c**). This suggests that PRL monomers can readily bind GAL monomers, possibly contributing to their initial interaction that eventually drives the synergistic aggregation and amyloid formation.

**Fig 4:**
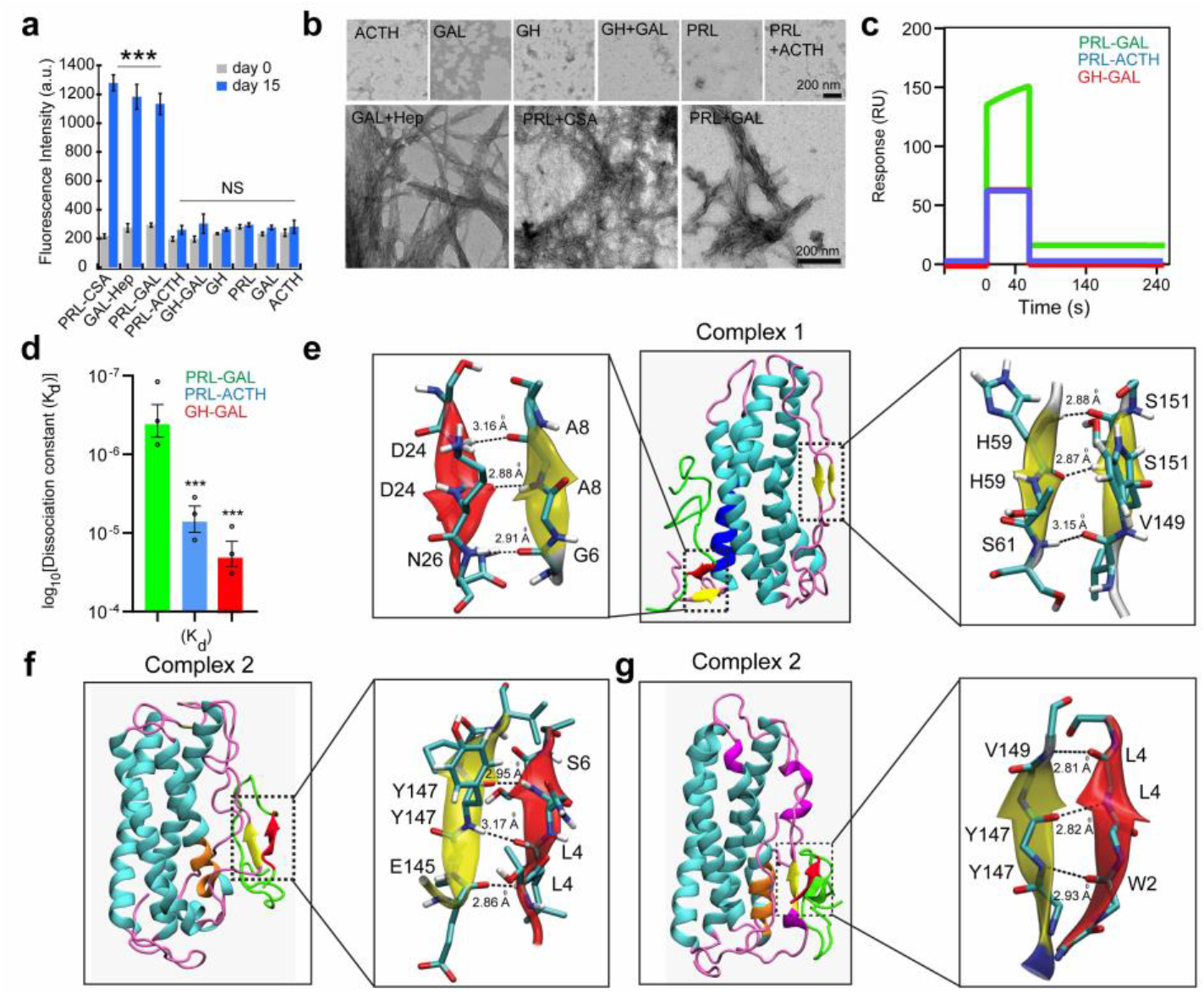
Specific interaction drives co-aggregation of PRL and GAL: **a**. Comparative ThT fluorescence showing amyloid formation by different pairs of hormones at days 0 and 15. PRL-CSA, GAL-Hep, PRL-GAL showed the highest ThT fluorescence signals after 15 days of incubation. Values represent mean ± SEM for n=3 independent experiments. The statistical significance is calculated between day 0 and day 15 for each sample using a t-test. **b**. The morphology observed under TEM for various hormones and the mixture of hormone samples are shown (after 15 days of incubation). Amorphous structures are seen for PRL-ACTH and GAL-GH; whereas PRL-GAL, PRL-CSA, GAL-Hep showed fibrillar morphology similar to amyloids. The experiment is performed three times with similar observations. **c**. Surface Plasmon Resonance (SPR) spectra showing strong binding of PRL on immobilized GAL compared to other pairs of hormones. **d**. The dissociation constant (K_d_) of PRL to GAL showing strong interaction between PRL and GAL for their co-aggregation and co-storage. The experiments are performed three times with similar results. Values represent mean ± SEM (***p ≤ 0.001, *p ≤ 0.05). **e**. Snapshot from *in silico* analysis (MD simulation) of PRL-GAL complex-1 using GROMOS 53a6 force field (when GAL is docked near residues 18-28 of PRL). **f**. Snapshot showing MD simulation of PRL-GAL complex-2 (when GAL is docked near residues 80-88 of PRL). Complex-1 induced the formation of an antiparallel β-sheet at the PRL-GAL interface (6-8 PRL & 24-26 GAL) and also an intra-molecular parallel β-sheet in PRL itself (59-61 PRL & 149-151 PRL). Complex-2 shows the formation of a parallel β-sheet constituted by the β-strand from PRL and GAL (145-147 PRL and 4-6 GAL). **g**. Snapshot showing MD simulation of complex-2 using amber ff99SB force field shows the appearance of parallel β-sheet at 147-149 residue of PRL and 2-4 residue of GAL. The snap-shot of complex-1 is included in **Supplementary Figure 11**.

To understand the mechanism of PRL and GAL interactions at the atomic level, docking, and molecular dynamics (MD) simulation studies were performed. GAL was docked at two different regions of PRL that showed high TANGO score (**Fig. 1b,c**). Thus, two sets of PRL-GAL complexes were generated (***a***) ***Set 1:*** GAL was docked near residues 18-28 of PRL and ***(b) Set 2:*** GAL was docked near residues 80-88 of PRL. The lowest energy docked complexes from each set, named complex-1 and complex-2 respectively, were obtained (**Supplementary Figure 11)**. Both these complexes were then subjected to independent 250ns long MD simulations to examine their stability. Since the choice of force fields may play an important role in the MD simulation results^62^, we performed MD simulations with two different force fields - GROMOS 53a6 force field^63^ and Amber ff99SB force field^64^. As controls, we have also simulated individual PRL and GAL proteins. We observed that the PRL and GAL alone did not show any noticeable structural changes (**Supplementary Figure 11**) during simulation time.

On the contrary, the PRL-GAL complexes exhibited significant conformational changes upon binding to each other, in both complexes (**Fig 4e-g)**. The respective structures of complex-1 and complex-2 at the end of the MD simulation using GROMOS force field showed the structural transition from unstructured region to β-strand in both PRL and GAL with the emergence of a parallel or anti-parallel β-sheet at the protein-protein interface (**Fig 4e,f**). The interaction of GAL in complex-1 induced the formation of an antiparallel β-sheet at the PRL-GAL interface (residues 6-8 of PRL, residues 24-26 in GAL) and also an intra-molecular parallel β-sheet in PRL itself (PRL residues 149-151 and 58-60) (**Fig 4e)**. In complex-2, GAL induced the formation of parallel β-sheet at PRL-GAL interface (residues 145-147 of PRL, residues 4-6 of GAL) **(Fig 4f)**. From the MD simulations using the Amber force field, however, the complex-1 did not show any notable structural changes, except that the terminal loops in PRL wrap around the existing secondary structures for higher stability **(Supplementary Fig 11)**. However, the PRL-GAL interactions in complex-2 resemble very well with the results from the GROMOS force field exhibiting a parallel β-sheet constituted by the β-strand from PRL and GAL proteins **(Fig 4g)**. These results corroborate very well with our experimental data that suggested the formation of amyloids when PRL and GAL were co-aggregated. The residues involved in the formation of this β-sheet in complex-2 were PRL residues 147-149 and GAL residues 2-4 in the Amber ff99SB force field (**Fig 4g**). Thus, irrespective of the force field used, our MD simulation results convincingly show that the co-aggregation of PRL and GAL induce the formation of β-sheet at the protein interface.

### Release of functional PRL and GAL from PRL-GAL amyloids

Protein/peptide misfolding and aggregation are known to lead to irreversible amyloid formation, which is stable and not readily disassemble to monomer. However, many studies recently showed the release of monomers and oligomers from disease-associated amyloids^65,66^. In contrast, amyloid formation related to SG biogenesis is reversible and should be able to release functional monomers in the extracellular space for their function^13,14,17^. To address whether any functional advantage of co-aggregation over homotypic aggregation by PRL and GAL, we determined the relative monomer release capability of PRL, GAL, and PRL-GAL co-amyloids, preformed fibrils of PRL (in the presence of CSA), GAL (in the presence of Hep) using dialysis method^13,14,17^. The concentration of released monomers (if any) in the dialysate was measured by UV-Vis spectroscopy at different time points. Intriguingly, we found that PRL-CSA, GAL-Hep, and PRL-GAL amyloids could indeed release monomers with time (**Fig 5a**). Interestingly, the amyloid fibrils of PRL and GAL formed in presence of GAGs (CSA and Hep, respectively) released monomeric hormones in a slow and sustained manner upon dilution in 10 mM Tris-HCl, pH 7.4 (**Fig 5a**).

**Fig 5:**
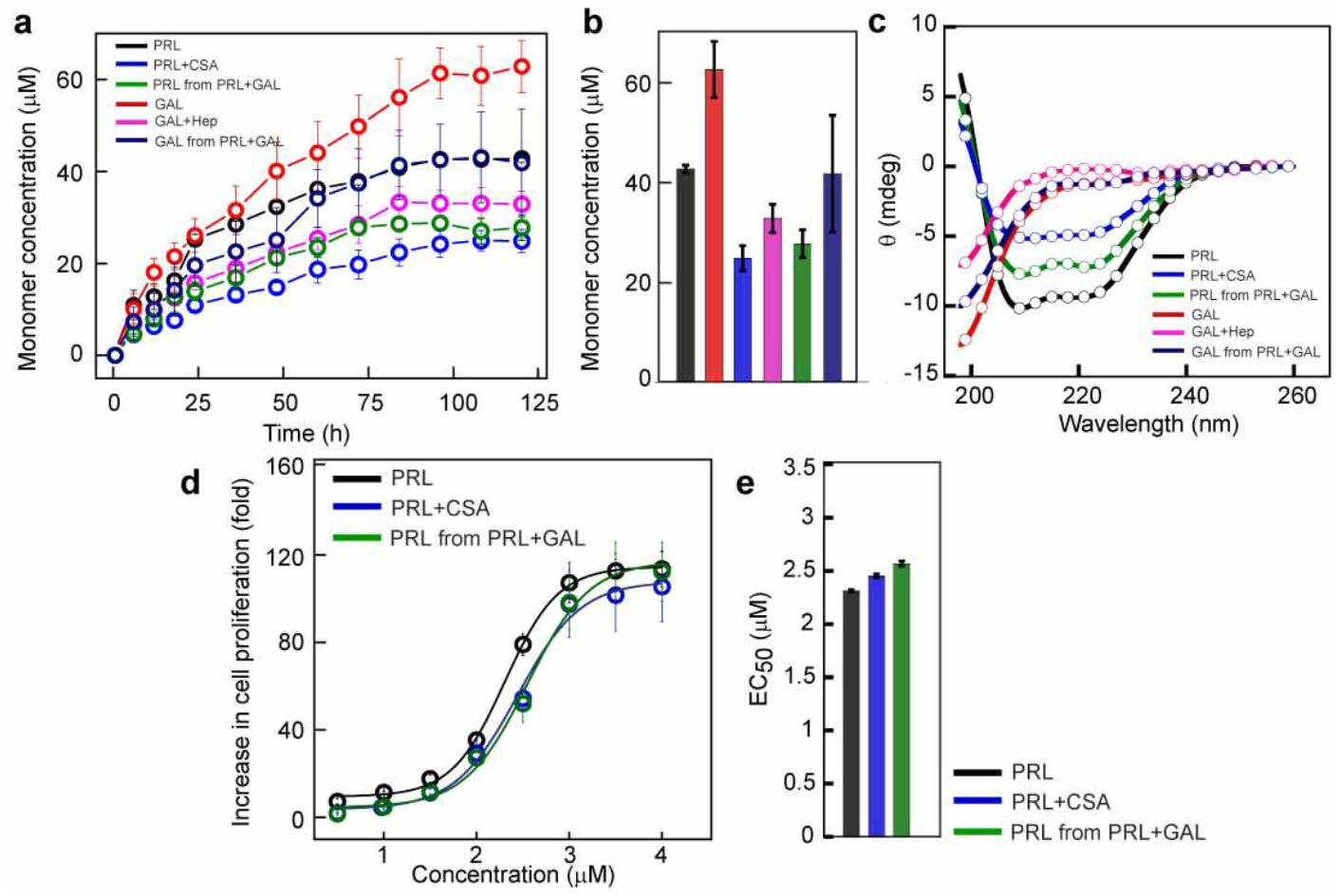
Monomer release from PRL and GAL amyloid. **a**. The kinetics of monomer release from various amyloids showing the continuous release of monomeric hormones. The experiment is performed three times with similar results. Values represent mean ± SEM for n=3 independent experiments. **b**. Saturation concentrations of different released monomers from fibrils along with the monomeric controls are shown. Values represent mean ± SEM for n=3 independent experiments. **c**. The secondary structure of released monomers showing their corresponding native secondary structures as confirmed by the CD. **d**. Nb2 cell proliferation study showing biological activity of released PRL from either PRL+CSA or PRL+GAL fibrils. Freshly dissolved protein was used as a control. Values represent mean ± SEM for n=3 independent experiments. **e**. EC_50_ values showing the released PRL monomers have similar bioactivity compared to freshly dissolved monomeric PRL. Values represent mean ± SEM for n=3 independent experiments.

However, the release of monomeric PRL and GAL hormones from the co-aggregated PRL-GAL fibril was faster compared to the PRL and GAL released from their GAGs mediated fibrils. This was confirmed by their release profile as well as saturation concentrations (**Fig 5a-b**). CD spectroscopy of the dialysate showed that the released PRL monomer retained its native conformation (**Fig 5c**).

Next, we performed the cell proliferation assay with the released PRL monomer to check for the retention of bioactivity of PRL. We used the Nb2 cell line for this study^67^. Nb2 cells are rat lymphoma cells with significant expression of PRL receptors on the cell surface^67,68^. These cells require PRL for proliferation or mitogenesis^68,69^. We observed that released PRL monomers obtained from PRL-CSA fibrils and PRL-GAL fibrils were functional as they can induce cell proliferation in a dose/concentration-dependent manner (**Fig 5d**). The EC_50_ values of the PRL released from PRL fibrils are similar as of freshly prepared PRL monomer, suggesting no difference in their functionality (**Fig 5e**).

## Discussion

Amyloids are ordered protein aggregates comprised of cross-β-sheet motifs where β-sheets are parallel, and individual β-strands are perpendicular to the fibril axis^47,70^. Despite their association with diseases, amyloids are also known to be involved in the native functions of host organisms including mammals^71,72^. Interestingly, the synergistic amyloid formation through co-aggregation and cross seeding by heterologous proteins/peptides is evident in the disease-associated amyloid formation such as α-Synuclein-Tau^73,74^, α-Synuclein-amyloid-β^28,75^. However, co-aggregation and cross seeding associated with functional amyloids are still elusive although this is of high functional relevance as many hormones are co-stored and co-released from secretory granules. Here, we explored the synergistic aggregation and amyloid formation by two human hormones prolactin (PRL) and galanin (GAL), with relevance to SG formation. Although 23 kDa PRL^41,42^ with mostly helical structure have no resemblance of sequence, length, and structure with unstructured 3.1 kDa GAL^43,44^, it was suggested that both of these hormones are co-stored in lactotrophs of anterior pituitary female rats^38^ and their release are also modulated by the same secretagogues^35,36,40^. Their co-storage and co-release suggest that they might aggregate together to form amyloid within the same secretory granules for storage. Indeed, the immunofluorescence study with anterior pituitary tissue showed not only both PRL and GAL do colocalize together but are also in the amyloid form as suggested by OC and ThioS staining (**Fig 1** and **Supplementary Fig 1**). Interestingly when PRL and GAL were co-incubated with a 1:1 or higher molar ratio *in vitro*, they co-aggregated synergistically to form amyloid fibrils in conditions similar to their storage in SGs without the requirement of any helper^76,77^ molecules.

PRL and GAL co-aggregation can happen where both PRL and GAL can promote the aggregation of each other but also form individual homotypic fibrils (**Fig 2**). Further, one of them might have the conformational advantage to form amyloids and other proteins/peptides can simply adhere to it (**Fig 2**). Also, PRL-GAL can be incorporated within the same fibrils or filaments (**Fig 2**). When immuno-EM was done with the secondary antibodies of different-sized gold nanoparticles, we observed both PRL and GAL are present within the same fibrils suggesting they might interact together to form hybrid fibrils (**Fig 2**). The incorporation of heterogenous protein to make hybrid fibrils is also shown in α-Synuclein (α-syn) and Tau and other fibril formtion^28,73-75^. The co-aggregation without GAGs where the individual hormone is unable to form amyloids further supports that in GAGs deficient condition, PRL and GAL might more prefer to form mixed amyloid for their storage. However, their co-aggregation does not preclude the possibility of individual storage of each hormone in the different SGs as both PRL and GAL formed amyloids in presence of respective helper GAGs molecules. Individual hormone aggregation can be seeded with their respective fibrils seeds as shown for PRL and GAL even in absence of GAGs, supporting the fact that amyloid fibrils formation might be very productive and formed autocatalytic manner inside the secretory cells.

Most often homologous seeding occurs for amyloid aggregation and decreases the lag time for aggregation^20,21^. Heterogeneous seeds also provide template/surface to protein/peptide for their aggregation, but not always cross seeding happens as barriers exist between two different protein/peptides^25,52,53^. It was shown that sequence similarity between two amyloidogenic proteins is crucial for cross seeding capability^78^. However, the ability of cross seeding also depends on the conformation of the seed and its compatibility with conformation or sequence of the monomeric protein, which creates different cross seeding barrier^25,51,79^ (also species barrier for prion diseases).

Our data showed PRL amyloid fibrils can cross-seed GAL monomers to form amyloid fibrils. Strikingly, preformed GAL fibrils were not able to cross seed PRL monomers suggesting specificity and regulation in functional amyloid formation (**Fig 3**). Such unidirectional, as well as bidirectional seeding between amyloid proteins, are also evident. For example, cross seeding between Aβ_42_ and hIAPP, where the Aβ_42_ seeds can cross seed the hIAPP to promote aggregation but hIAPP seed was unable to induce aggregation to Aβ_42_ monomers^80^, suggesting the unidirectional cross seeding. In contrast, α-syn seeds speed up the aggregation of Tau and Tau seeds also accelerated the aggregation of α-syn suggesting the bidirectional cross seeding^73,74^.

Since PRL has less propensity to form amyloid due to higher structural stability, GAL seeds are not able to provide neither surface nor structural compatibility for PRL for cross seeding. This might suggest that tight regulation of co-species aggregation ensures the right amount of storage of PRL and GAL for secretory storage with proper ratio. Moreover, GAL is a highly disordered peptide, which might easily bind to the PRL amyloid surface nonspecifically and increase its stability and aggregates. Further PRL amyloid-prone sequence could also be sequestered inside the helical structure, which might not be compatible with either surface and/or amyloid core structure of GAL to mediate the cross-seeding. Previously it was shown that cross seeding of K18 and K19 of Tau isoforms, K19 fibrils can cross seed K18 through the catalytic motif of R3, whereas K18 fibrils with the catalytic center as R2 is unable to seed K19^81^ (lacking R2 motif). The difference in seeding could also be due to the relative tendency of amyloid fibril formation by GAL and PRL. We propose that PRL fibrils seeding GAL aggregation is not due to elongation and fragmentation (which require specific interaction) as PRL cross-β spine might not be assessable for GAL. We previously proposed that a small segment of GH (structurally similar to PRL) might engage for fibrils formation^14^ and other structural domains might be surrounded to the cross-β spine as proposed for RNaseA fibrils model^82^. We believe due to the unstructured peptide of GAL, it could easily adhere to the surface of PRL amyloid seeds and proceeds for surface-mediated secondary nucleation. This is further supported as the structure of PRL seeded GAL fibril showed similar fibrils morphology and secondary structure as of GAL fibrils formed alone (**Fig 3**). This is a typical property of surface-mediated secondary nucleation that produce fibrils^56^. Further the PRL seeding to GAL fibril formation was more efficient as compared to homo-seeding of GAL further support surface mediated secondary nucleation^53,56^ (**Fig 3**).

We hypothesized that specific interaction between GAL and PRL synergistically facilitates their aggregation into amyloid. This is further evident from the SPR analysis showing strong interaction and direct binding between the monomeric forms of PRL and GAL in contrast to the other hormone pair of PRL-ACTH or GH-GAL (**Fig 4**). This PRL-GAL interaction and conformation transition are further supported using *in silico* study, which showed interaction of the PRL N-terminal loop and GAL, where GAL promotes the conformational transition of the N-terminus of PRL into β structure (**Fig 4**). The interaction of PRL and GAL is mandatory for their aggregation, as amyloids were not formed when PRL and GAL were incubated alone.

The observed co-aggregation and cross-seeding may not only give the advantage for both PRL and GAL to be stored as amyloids (and nonrequirement of helper molecules) but also may help for their subsequent release after secretion. This is supported by the fact that the monomer release from PRL-CSA amyloid and GAL-Hep amyloid was much slower compared to the release of mixed amyloid of PRL-GAL (**Fig 5**). Further, our study suggests that nature has optimized hormone storage in such a way that one hormone can be stored in different ways in different secretory granules and the release of these hormones also depends on the type of amyloid aggregate, which is stored in the SG (**Fig 6**). The present study suggests that contrary to disease-associated co-aggregation, which promotes the spread of the disease, co-aggregation of hormones is specific and functional in the SGs. Co-aggregation and amyloid formation of these structurally dissimilar hormones indicate the relevance of amyloid as a crucial aspect of cellular sorting and storage in SG biogenesis.

**Figure 6:**
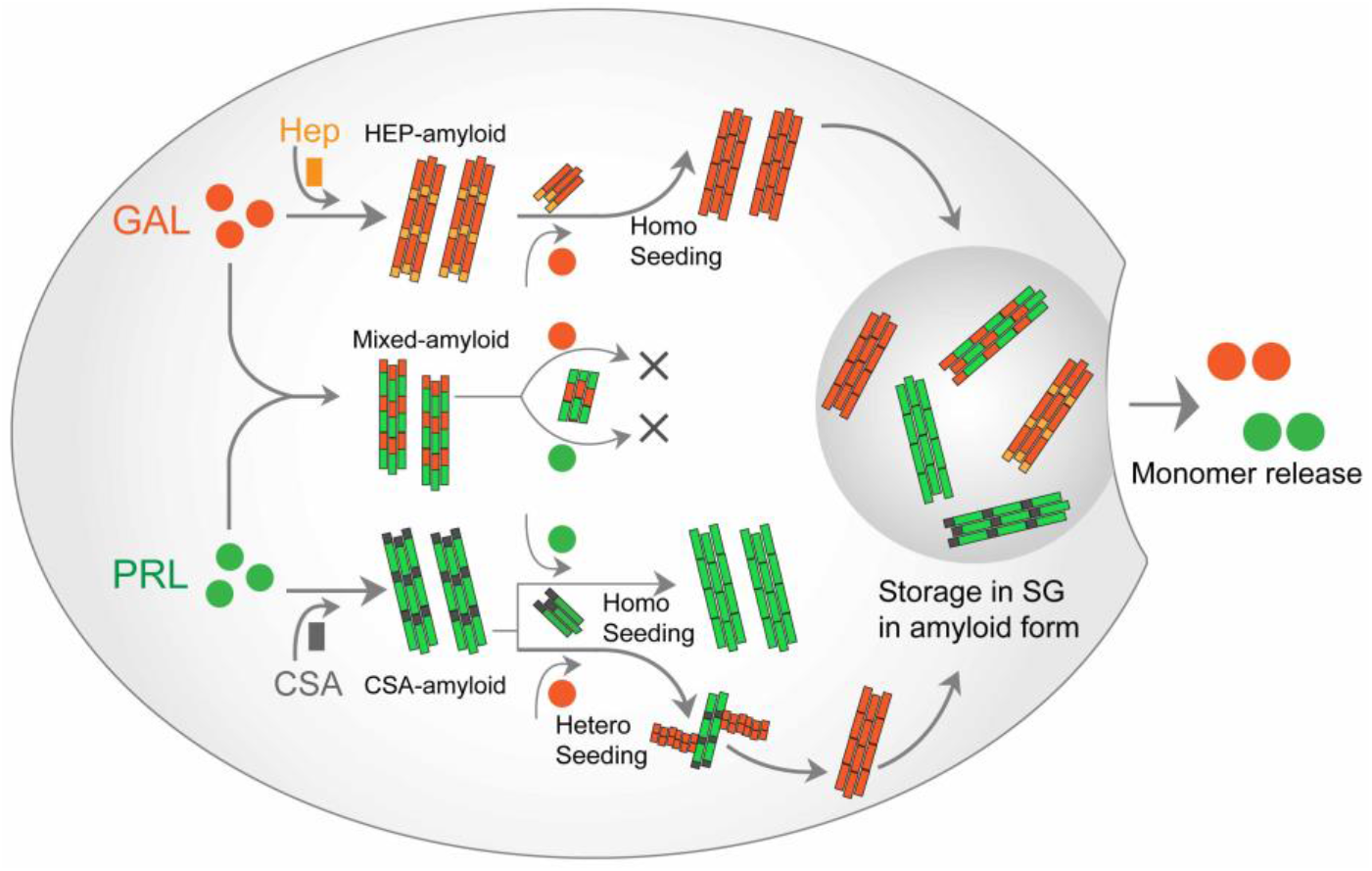
PRL-GAL homo and hetero amyloid life cycle for secretory granule. PRL and GAL form the amyloid fibrils in the presence of specific GAGs (CSA and Hep, respectively), which can be auto-catalytically amplified by their respective seeding with preformed fibrils. This seeding however does not require any GAGs. PRL-GAL also synergistically co-aggregate to form hybrid amyloid fibrils, which are not capable of seeding either to PRL or GAL. These amyloid fibril species can together or individually reconstitute the SGs of PRL-GA storage, which can release functional PRL and GAL into the extracellular space.

## Materials and Methods

### Chemicals and Reagents

The peptide GAL was purchased from USV Limited (Mumbai, India) by custom synthesis and other chemicals were obtained from Sigma Chemicals or other sources with the highest purity available.

### Expression and purification of human Prolactin (PRL)

PRL was expressed as per the protocol reported^83^ with little modification. The human PRL plasmid was obtained as a kind gift from Prof. Dannies and Prof. Hodsdon from Yale University. BL21 (DE3) *E. coli* cells were transformed with the hPRL gene encoded in the pT7L plasmid and were made to grow in terrific broth (TB) followed by IPTG induction for 4 hours. After harvesting the cells at 8000 rpm for 20 minutes, it was dissolved in 20 mM Tris-HCl, pH 8.0 (with added protease inhibitor cocktail, Roche). After that, the cells were sonicated (2 seconds on, 1 second off at 50% amplitude) for 20 minutes to complete cell-lysis, which was then centrifuged at 15000 rpm for half an hour to recover the inclusion bodies (IB). The cell pellet containing IBs was subsequently washed two times with 0.5% triton-X and then it was dissolved in 8 (M) urea with 2% (v/v) β-mercaptoethanol. The solution was then dialyzed against 20 mM Tris-HCl, pH 8.0 so that the PRL protein refolds to its native state. After dialysis, the protein solution was again centrifuged at 15000 rpm for 1 hour and was loaded in an anion exchange column (Resource Q, GE healthcare) through an AKTA purifier FPLC system (Cytiva). The protein was eluted through 1(M) NaCl gradient and subsequently lyophilized after snap-freezing in liquid nitrogen. Size exclusion profile (SEC) suggested the protein is monomeric in nature and the purity of the protein is further checked by SDS-PAGE and MALDI-TOF spectrometry. CD spectroscopy was also performed to confirm that the purified PRL has refolded to its native helical conformation.

### Aggregation of PRL and GAL in the presence of GAGs

PRL was dissolved in MQ water and buffer exchanged to 20 mM phosphate buffer with 100 mM NaCl, pH 6.0, 0.01% sodium azide using 10 kDa mini dialysis units (Thermo Scientific Slide-A-Lyzer). 5 mM solution of CSA (Sigma, USA) was prepared in the same buffer, and appropriately mixed with PRL to obtain an ultimate concentration of 400 μM for both PRL and CSA. Similarly, for GAL aggregation in the presence of Hep, GAL peptide was dissolved in 20 mM phosphate buffer containing 100 mM NaCl, pH 6.0, 0.01% sodium azide to obtain a concentration of 500 μM. Hep solution from a stock of 5 mM (made in the same buffer) was then mixed with GAL solution to obtain an ultimate concentration of 400 μM for both GAL and Hep. These tubes containing PRL and GAL of various mixtures were kept into an Echo Thermmodel RT11 rotating mixture with a speed of 50 rpm for 15 days inside a 37°C incubator. As a control, 400 μM PRL and GAL alone in the same buffer, was also incubated in a similar condition. The secondary structural transition was monitored by CD spectroscopy and amyloid formation by ThT binding assay at various time points. Finally, Congo red (CR) binding studies and TEM imaging was used to confirm amyloid fibril formation.

### Co-aggregation study of PRL and GAL

For the co-aggregation study, PRL was dissolved in Milli-Q water and buffer exchanged to 20 mM phosphate buffer with 100 mM NaCl, pH 6.0, 0.01% sodium azide using 10 kDa mini dialysis units (final concentration was 800 μM) (Thermo Scientific Slide-A-Lyzer). GAL was also dissolved in the same buffer to obtain an 800 µM solution. After that, each of the solutions was mixed to obtain 400 µM of PRL-GAL mix and was incubated with slight agitation at 37°C for two weeks. 400 µM of each PRL and GAL was incubated alone as controls. For co-aggregation of PRL and GAL with other hormones, separate solutions of PRL, GAL, GH, and ACTH were freshly dissolved and prepared in identical solution condition as above. 200 µl of each solution were mixed to obtain 400 µM each of PRL-ACTH and GAL-GH mixture and was incubated with slight agitation at 37 °C for two weeks. 400 µM of each of PRL, GAL, GH and ACTH were also incubated alone as controls. The secondary structural transition and amyloid formation of the incubated solutions were monitored by CD spectroscopy and ThT binding assay during various time points. After 15 days of incubation, the morphology of the incubated samples was analyzed by TEM.

### Circular Dichroism spectroscopy (CD)

For CD measurement, protein/peptide aliquots were diluted in 20 mM phosphate buffer with 100 mM NaCl, pH 6.0, 0.01% sodium azide to 200 μl and the final concentration protein/peptide was 10 μM. CD spectra were taken using a JASCO 810 instrument where the sample was loaded in a quartz cell of 0.1 cm path length (Hellma, Forest Hills, NY). Spectra were collected at 198-260 nm wavelength (far UV) at 25°C. Raw data was processed by smoothening, as per the manufacturer’s instructions. Three independent experiments were performed with each sample.

### Thioflavin T (ThT) Binding Assay

To measure ThT binding, PRL and GAL solutions were diluted in the same buffer into 200 μl such that final concentration of each sample was 10 μM. 4 μl of 1 mM ThT prepared in 20 mM Tris-HCl buffer, pH 8.0 was added into each sample. ThT fluorescence was probed after the immediate addition of ThT. The fluorescence experiment was carried out in Shimadzu RF5301 PC, with excitation wavelength at 450 nm and emission wavelength from 460-500 nm. For measuring both excitation and emission, the slit width was kept at 5 nm. Three independent experiments were performed for each sample.

The lag time (t_lag_) was calculated as per the published protocol^84^:

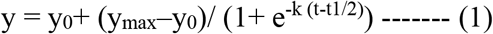

here y is the ThT fluorescence at any particular time point, y_max_is the maximum ThT fluorescence observed and y_0_ is the ThT fluorescence at t_0_ (initial time) and t_lag_ was defined by

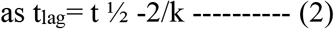

### Congo red (CR) binding

A 5 μl aliquot of protein/peptide sample was added into 80 μl of 5 mM potassium phosphate buffer containing 10% ethanol. 100 μM CR solutions were prepared in 5mM phosphate (containing 10% ethanol) and 15 μl of the solution was added to the sample. After incubating for 15 min in dark, absorption spectra were taken from 300-700 nm (JASCO V-650 spectrophotometer). For control CR solution without protein was also measured. Three independent experiments were performed for each sample.

### CR birefringence study

Protein fibrils were obtained by ultracentrifuging the fibril solution at 95,000 rpm for 1 hour followed by washing with Milli-Q water. The fibrils were mixed in 100 μl of alkaline sodium chloride solution for 20 min with vortexing, to ensure uniform mixing of all fibrils in solution. The mixture was further centrifuged and pellet fractions were stained with alkaline CR solution for 20 minutes with vortexing. After that, mixtures were again centrifuged at 95,000 rpm for 1 hour, and pellets were washed two times by 500 μl of 20% ethanol. The pellets were then resuspended in PBS and spotted onto glass slides and subjected to air-drying at room temperature. The slides were observed using a microscope (Olympus SZ61 stereo zoom) attached with two polarizers and a camera.

### Immunoelectron microscopy

10 µl of PRL-GAL or PRL-CSA fibril was spotted onto the TEM grid. 10 µl of rabbit anti-PRL antibody (1:10) and/or mouse anti-GAL antibody (1:10) in PBS was added to the fibrils and was incubated for 1 h. The excess antibody was removed using filter paper. The grid was subsequently washed thrice with autoclaved MQ water. Anti-mouse secondary antibody conjugated with 10 nm gold particles (1:200) and/or anti-rabbit secondary antibody conjugated with 5 nm gold particles (1:200) was added to the grid and incubated for 30 min. The grid was then washed thrice with MQ water, followed by staining with 1% uranyl formate for 5 min and it was imaged using TEM (CM200, Netherland), and analyzed using KEEN view software.

### Transmission electron microscopy (EM)

The protein/peptide sample was diluted in Milli-Q water to ∼60 μM. Then, the samples were spotted on a carbon-coated, glow-discharged Formvar grid (Electron Microscopy Sciences, Fort Washington, PA) and were kept for incubation for 5 min. The grids were further washed with Milli-Q water and were stained with a 1 % (w/v) uranyl formate solution. TEM imaging was done using FEITecnai G^2^ 12 electron microscope at either 120 kV or 200 kV with nominal magnifications in the range of 26,000–60,000. Images were collected by using the SIS Megaview III imaging system. Independent experiments were carried out thrice for each sample.

### X-ray fibril diffraction

PRL/GAL fibrils were isolated by ultracentrifugation as mentioned earlier and were loaded into a clean 0.7 mm capillary. The samples in capillary were dried overnight under vacuum. The whole capillary with dried protein was placed in the path of X-ray beam. The dried film of protein was placed in an X-ray beam at 200 K for 120 s exposure. The resulting images were collected using a Rigaku R-Axis IV++ detector (Rigaku, Japan) kept on a rotating anode. The distance between the sample to the detector was 200 mm and the image files were analyzed and processed using Adxv software.

### FTIR spectroscopy

For FTIR spectroscopy, isolated fibrils or monomers were spotted onto a thin KBr pellet and were subjected to dry under an IR lamp. Then the spectrum was collected using a Bruker VERTEX 80 spectrometer attached with a DTGS detector at the frequency range of 1800-1500 cm^-1^, corresponding to the amide I stretching frequency, with a resolution limit of 4 cm^-1^. The recorded spectrum was deconvoluted at the frequency range 1700-1600 cm^-1^, using Fourier Self Deconvolution (FSD) method and the deconvoluted spectrum was fitted using the Lorentzian curve fitting method using OPUS-65 software (Bruker, Germany) according to the manufacturer’s instructions. Independent sets were performed thrice for each sample.

### Seeding and cross-seeding by different fibril-seed

Amyloid formation by PRL/GAL was confirmed by ThT binding and TEM imaging. After that, various fibrils were collected seperately via ultra-centrifugation, and each fibril sample was suspended in 20 mM phosphate buffer, pH 6.0, 100 mM NaCl, 0.01% sodium azide. These fibrils were then subjected to sonication (03 seconds“on” and 01 second “off” at 20% amplitude) for 10 minutes to obtain preformed fibril seeds, which were mixed in the respective monomeric protein (1%, 2% and 5% (v/v)) for homo seeding or to the other protein for cross seeding. The time-dependent aggregation was probed by ThT binding and CD spectroscopy.

### Surface plasmon resonance (SPR) spectroscopy analysis of PRL-GAL interaction

GAL was immobilized on to Biacore CM5 sensor chip (GE Healthcare) via amine coupling. To do that, 300 µg/ml of GAL solution was made in 50 mM sodium acetate buffer, pH 5, and was injected at a rate of 10 µl/min for 720 s to achieve a response unit (RU) of 1252 RU. PRL protein was dissolved in 20 mM phosphate buffer containing 100 mM NaCl, pH 6.0 to obtain 1 mM stock solution and was used for preparing 7.8, 15.6, 31.2, 62.4, 125, 250, and 500µM dilutions. These solutions were then injected over the immobilized GAL at a flow rate of 45 µl/min for 60 s. The dissociation was initiated at a flow rate of 30µl/min, and the signal was recorded for 300 s. The chip was further regenerated using a 10mMNaOH solution. ACTH and GH protein were used to examine the binding with PRL and GAL, respectively using a CM5 chip. For immobilization, 500 µg/ml of ACTH solution was made in 50 mM sodium acetate buffer (pH4.5). For PRL immobilization, 2 mg/ml of PRL protein in 50 mM sodium acetate buffer (pH 4) was used. A similar range of concentrations was used for ACTH and GH as mentioned before. All the binding experiments were performed at 37 °C. A blank run (without analyte) was made to be subtracted from the sample response in addition to the reference surface, to rectify any instrument noise due to injections. The RU values obtained for each experiment were fitted using a steady-state, 2-step model and the relative dissociation constant (K_d_) was calculated using Biacore software.

### *In silico* study of PRL and GAL interaction and co-aggregation

Protein-protein docking and all-atom molecular dynamics (MD) simulations were used to probe the co-aggregation propensities (if any) and resulting secondary structural transitions in PRL and GAL molecules. For this, the initial structure of PRL was obtained from PDB ID: 1RW5^41^. Since there was no entry for GAL structure in PDB, the GAL structure was built from its primary amino acid sequence and energy minimized. Subsequently, the minimized structure was equilibrated and simulated for 300ns. Initially, GAL was docked at two amyloidogenic regions of PRL (residues 18-28) (set-1) and (residues 80-88) (set-2), which were predicted by TANGO^45^. The docking was performed by the protein-protein docking program, HADDOCK^85^. The lowest energy complexes were extracted from each set, denoted as complex-1 from set-1 and complex-2 from set-2. These two complexes were used as starting structures for the MD simulations. The first sets of simulations were performed using the AMBER16 package with the Amberff99SB force field. The LEAP module of AMBER16 was used to add the hydrogen for the heavy atoms. The complexes were then energy minimized for 2000 steps using the steepest descent and conjugate gradient algorithms. Subsequently, the structures were hydrated in a cubic periodic box extending 9Å outside the protein-protein complex on all sides with explicit water molecules. The three-site TIP3P model was chosen to describe the water molecules. The charge of each system was neutralized by placing Na^+^ ions randomly in the simulation boxes. The systems were again minimized to prevent any bad contacts formed due to the solvation. All the systems were then equilibrated for 500ps in NVT ensemble at 300K followed by 1 ns in NPT ensemble at 1 atm of pressure. After the density and potential energy of the systems had converged, each complex was subjected to 300 ns of the production run. To further validate our simulations, we performed a new set of simulations using different force fields. Each complex was subjected to a 250ns simulation using the GROMOS 53a6 force field. The new sets of simulations were performed using the GROMACS package following the above-mentioned protocol. VMD tool was used for visual analysis of the trajectories.

### Monomer release assay

Amyloid fibrils of PRL and GAL formed in the presence of GAGs, and co-aggregated fibrils of PRL-GAL were harvested by ultracentrifugation at 90,000 rpm for 1 h. The concentration of the soluble fraction (supernatant) was calculated using the absorbance at 280 nm (Jasco V-650) and was used to determine the concentration of the pelleted fibrils. 100 μl re-dissolved pellet of 400 μM concentration was chosen to examine the monomer release study using the experimental setup, which has been reported previously^14,86^. 400 μM PRL and GAL solutions incubated for 15 days were used as a monomer control. Briefly, For PRL monomer release, the pellet solutions (PRL, PRL-CSA, and PRL-GAL) were transferred into a modified PCR tube with a pierced hole in its cap, which was attached and sealed with a 50 kDa molecular weight cutoff membrane (Pierce, USA). This whole setup was then placed inside a 1ml cryotube (Nunc, Denmark) containing 500 μl of 10 mM Tris-HCl buffer (pH 7.4), 0.01% sodium azide. Meanwhile, to investigate the release of GAL monomers from the fibril of GAL-Hep, pellet solution was placed in a 10 kDa cutoff Slide-A-Lyzer mini dialysis unit system (Pierce, USA), which was positioned onto a 1 ml cryo-tube (Nunc, Denmark) containing 500 μl of Tris-HCl buffer (pH 7.4), 0.01% sodium azide. The tubes were then kept at 4°C to prevent evaporation of solutions. To calculate the concentration of the protein outside of the membrane, an aliquot of 100 μl of the solution was pipette from the buffer outside of the dialysis membrane at different times, absorbance was measured at 280 nm. For measuring the monomer release of PRL and GAL from PRL-GAL co-aggregates, 100 μl releasing medium was taken out, and the same volume of fresh buffer was added back as volume correction (subsequent concentration correction has been done in the data). The separated 100 μl releasing medium was passed through a 10 kDa centrifugal filter unit (Amicon Ultra, Millipore). After which, the GAL monomer was obtained as filtrate, and the PRL was recovered from the retentate by washing the reversed filter unit with 100 μl of 10 mM Tris, pH 7.4 as per the manufacturer’s protocol. In an identical experimental setup, the absorbance was measured from the buffer both inside and outside of the membrane. This control is kept to check whether degradation of the materials of the dialysis membrane is interfering with the assay. Each time, the released solution was returned after the spectra recording. Three independent experiments were performed for each sample.

### Cell proliferation assay using Nb2 cells

Nb2 cell line was grown in plastic culture flasks in RPMI medium supplemented with 10% heat-inactivated fetal bovine serum (FBS), 10 % horse serum (HS), and 1X antibiotic solution and incubated at 37°C in a humidified incubator containing 5% CO_2_ in the air. For proliferation assay, cells (∼ 10^5^/well) in a 96 well plate were seeded in RPMI medium in the presence of 1 % fetal bovine serum and 10 % HS and incubated for 24h to synchronize the cells at G0/G1 phase. After 24h, cells were treated with PRL monomer, monomer released from PRL-CSA and PRL-GAL at a dose range from 0.5 – 4 µM. The unrelated protein ovalbumin was used as the negative control. After incubation, cell proliferation was measured by MTT assay. To do so,10 µl of MTT solution (5mg/ml in PBS) was added to the cells and incubated for 4 h. Subsequently, 100 μl of SDS-DMF solution (50% DMF and 20% SDS, pH 4.75) was added for overnight incubation. The absorption value of the product was measured at 560 nm and 690 nm as a background absorbance using a Spectramax M2e microplate reader (Molecular Devices, USA). The fold increases in cell proliferation compared to untreated cells were plotted against the concentration of the sample administered.

### Immunofluorescence

Adult, female, Sprague-Dawley rats (200-250 g) taken for this study were maintained under the standard environmental conditions (12h: 12 h, light: darkness cycle, chow and water *ad libitum*) and Institutional Animal Ethical Committee (IAEC) at NISER, Bhubaneswar, India approved the experimental protocols. First, the animals were anesthetized with sodium pentobarbital and then perfused transcardially with 50 ml of 10 mM phosphate buffer saline (PBS) pH 7.4, followed by 50 ml 4% paraformaldehyde (PFA) in 100 mM phosphate buffer pH 7.4. The pituitary glands were dissected out and post-fixed in 4% PFA overnight at 4°C followed by immersion in paraffin. Thin sections of the various paraffin-embedded tissues were made using a microtome. The sections were deparaffinized and rehydrated in decreasing concentrations of xylene and increasing concentrations of ethanol. The sections were rinsed twice in distilled water, followed by enzymatic antigen retrieval using 0.05% Trypsin incubation at 37°C for 5 min. The sections were then washed with TBST buffer, pH 7.4 (Tris-buffered saline containing 0.1% tween-20) followed by treatment with 0.2% Triton X-100 for 10 min. The sections were blocked using TBST containing 2% BSA to prevent non-specific binding. PRL-GAL co-immuno-staining was performed using anti-PRL(Guinea pig polyclonal, from A. F. Parlow, National Hormone, and Pituitary Program, Harbor-ULCA Medical Center, Torrance, CA,1:1500) and anti-GAL primary antibody (Mouse, Abcam) (1:1500) overnight at 4°C. Further, co-staining of pituitary tissue amyloids and GAL or PRL was performed using amyloid-specific (OC) antibody (rabbit polyclonal, Abcam, 1:500) and with respective hormone antibodies. All of the primary antibody-stained tissue slices were incubated overnight at 4 °C in a humidified chamber. The sections were rinsed in TBST and further incubated with the secondary antibody of goat anti-mouse FITC (1:500) or goat anti-rabbit Alexa Fluor-647(1:500) (Life Technologies, Thermo Scientific, USA) or goat anti-Guinea pig Alexa Fluor 555(1:500)for 2 h at room temperature in a humid chamber. The sections were washed with TBST and mounted with a mounting medium. The sections were analyzed using a confocal microscope (Olympus IX81 combined with FV500) (Shinjuku, Tokyo, Japan) and images were recorded using a multi-channel image acquisition tool of Fluovision software (Zeiss, Oberkochen, Germany). For Thioflavin S (ThioS) staining, GAL and PRL staining was done using goat anti-mouse Alexa Fluor-555-conjugated secondary antibody (1:500 dilution) or goat anti-Guinea pig Alexa Fluor 555 respectively. The sections were then stained with 0.6% ThioS (Sigma-Aldrich) for 5 min in dark. The sections were washed with 50% ethanol for 2 min followed by TBST washing for 3 min. The slides were then mounted in 90% glycerol and 10% phosphate-buffered saline (PBS) containing 1% DABCO (1, 4-diazabicyclo-[2.2.2] octane, Sigma-Aldrich). The images and the sections were analyzed by multi-channel image acquisition tool of Fluovision software (Zeiss, Oberkochen, Germany) and Olympus FV-500 IX 81 confocal microscope (Shinjuku, Tokyo, Japan) respectively.

## Acknowledgments

Authors wish to acknowledge Prof. P. S. Dannies and Prof. M. E. Hodsdon, Yale School of Medicine, USA for the plasmid of prolactin. Srivastav Ranganathan and Prem Prakash for the PRL schematic drawing and Congo red birefringence study, respectively. We are also thankful to CRNTS and IRCC, IIT Bombay for FTIR, electron microscopy, protein crystallography facilities and SPR facility. Authors wish to acknowledge DBT (BT/PR9797/NNT/28/774/2014) Government of India, Wadhwani research centre for Bioengineering (WRCB) and DBT/Welcome Trust India Alliance Fellowship [RD/0119-DBTFL49-001] awarded to Shinjinee Sengupta for financial support.

## Author contributions

D.C., R.S.J and S.K.M designed the experiments. D.C., R.S.J., S.R., A.N., N.S., S.S., L.G., P.K., D.D., A.P., C.P., S.K., P.S., and S.S performed experiments. All authors analyzed the data. S.K.M, D.C., S.R. & R.S.J wrote the manuscript. All authors approved the final version of the manuscript.

## Data availability statement

The authors declare that all the data supporting the findings of this study are available within the paper and in supplementary information files. All the data analysis was performed using published tools and packages and has been cited in the paper and supplementary information text.

