## Supplementary Figures and Table for "Co-aggregation and secondary nucleation in the life cycle of human prolactin/galanin functional amyloids"

**This file includes:**

Supplementary Figures S1-S11
Supplementary Table 1

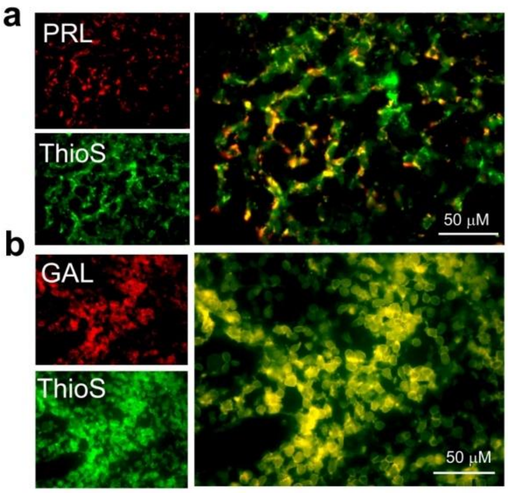


**Supplementary Fig 1**: **ThioS co-localization of PRL and GAL in female rat pituitary tissue.** Representative fluorescence microscopic images of female rat anterior pituitary tissue (after immunostaining) showing ThioS co-localization with both the hormones. The data indicate that PRL and GAL are stored as amyloid in secretory granules (SGs). **a.** PRL (red) and ThioS (green) shows co-localization (yellow). **b.** GAL (red) and ThioS (green) shows co-localization (yellow). The experiment is performed three times with similar observations.

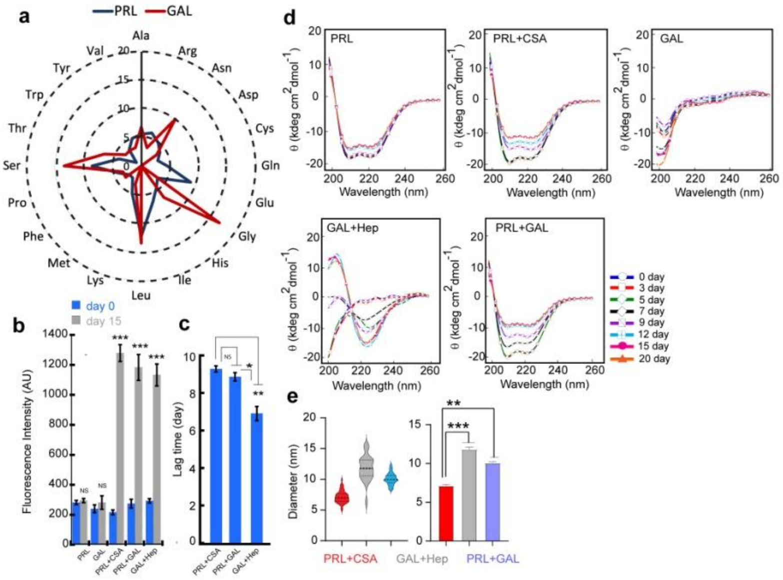

**Supplementary Figure 2: Conformational transition and aggregation by PRL and GAL**
**a.** Radar plot showing no significant sequence similarities between PRL and GAL. **b.** ThT fluorescence at day 0 and day 15 for PRL, GAL, GAL-Hep, PRL-GAL, and PRL-CSA is shown. The data showing a significant increase in ThT fluorescence for GAL-Hep, PRL-GAL & PRL-CSA after 15 days of incubation. PRL and GAL alone do not show any significant ThT binding even after 15 days of incubation. Values represent mean ± SEM for n=3 independent experiments. Statistical significance is calculated for each sample using paired Student's t-test. **c.** Lag time of aggregation derived from ThT aggregation kinetics of PRL-CSA, PRL-GAL, and GAL-Hep. Values represent mean ± SEM for n=3 independent experiments. The statistical significance(***$p \leq 0.001$, **$p \leq 0.01$, *$p \leq 0.05$) is calculated by one-way ANOVA followed by an SNK post hoc test with a 95% confidence interval **d.** CD spectra showing secondary structural changes during incubation. PRL-GAL and PRL-CSA show a decrease in helicity with time. GAL-Hep shows a structural transition from RC to the β-sheet structure during incubation. The experiment is performed three times with similar results. **e. (*Left panel*)** Median values of different fibril diameters are shown with violin plots. (***Right panel***) Average values of different fibril diameters are shown. Values represent mean ± SD for n=3 independent experiments. The statistical significance(***$p \leq 0.001$, **$p \leq 0.01$, *$p \leq 0.05$) is calculated by one-way ANOVA followed by an SNK post hoc test with a 95% confidence interval.

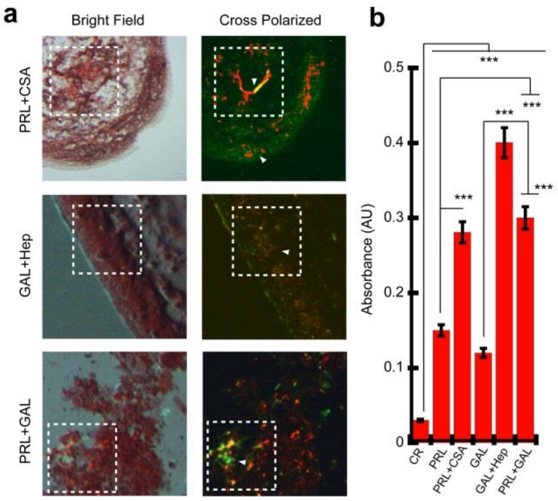


**Supplementary Fig 3: Congo red (CR) birefringence of hormone amyloids a.** Bright-field and cross-polarized images of the PRL-CSA, PRL-GAL, and GAL-Hep samples after 15 days of incubation are shown. The greenish-yellow CR birefringence under crossed polarized light indicates the presence of amyloid in all three samples. **b.** PRL-CSA, GAL-Hep, and PRL-GAL showing higher CR absorbance compared to PRL and/or GAL alone samples. Values represent mean ± SEM for n=3 independent experiments. The statistical significance (***$p \leq 0.001$) is calculated by one-way ANOVA followed by the SNK post hoc test with a 95% confidence interval.

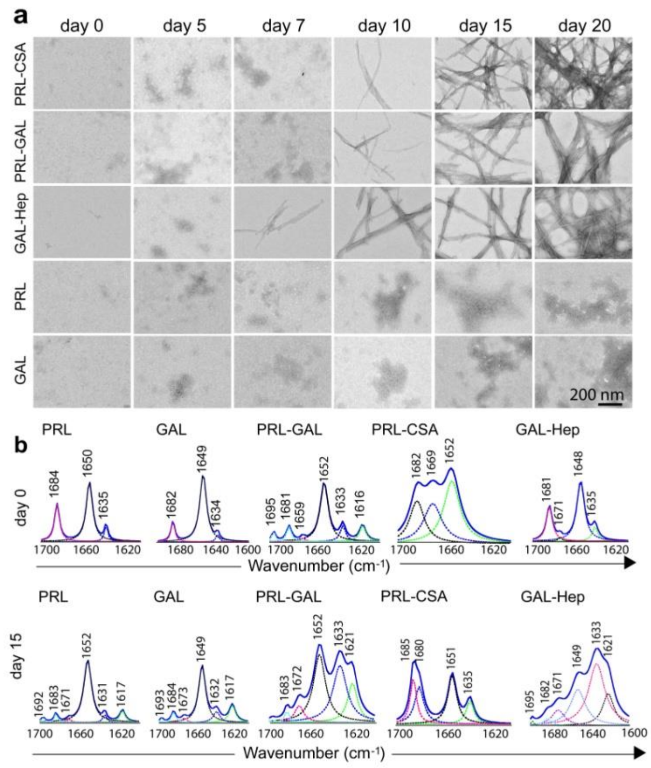


**Supplementary Fig 4: Time dependent amyloid fibril formation by hormones using TEM and FTIR study. a.** The morphology of the formation of the aggregates was monitored by TEM over the aggregation time-course. The data showing the appearance of fibrillar structure for GAL-Hep in a faster time point followed by PRL-GAL and PRL-CSA. Only PRL and/or GAL show amorphous morphology throughout the aggregation time-course. The experiment is performed three times with similar observations. Scale bar is 200 nm as shown. **b.** FTIR spectra of PRL-GAL, PRL-CSA & GAL-Hep along with PRL and GAL alone at day 0 and day 15 is shown. On day 0, PRL, PRL-CSA & PRL-GAL shows the major peak at 1651 $cm^{-1}$, which is characteristic of the α-helical structure. GAL & GAL-Hep shows the major peak at 1648 $cm^{-1}$, which is characteristic of random coil (RC) structure. At day 15, PRL-GAL shows two peaks at 1633 and 1621 $cm^{-1}$, representing β-sheet conformation. PRL-CSA shows a peak at 1636 $cm^{-1}$ and GAL-Hep shows peaks at 1621 & 1633 $cm^{-1}$ representing the β-sheet structure. The experiment is performed three times with similar observations.

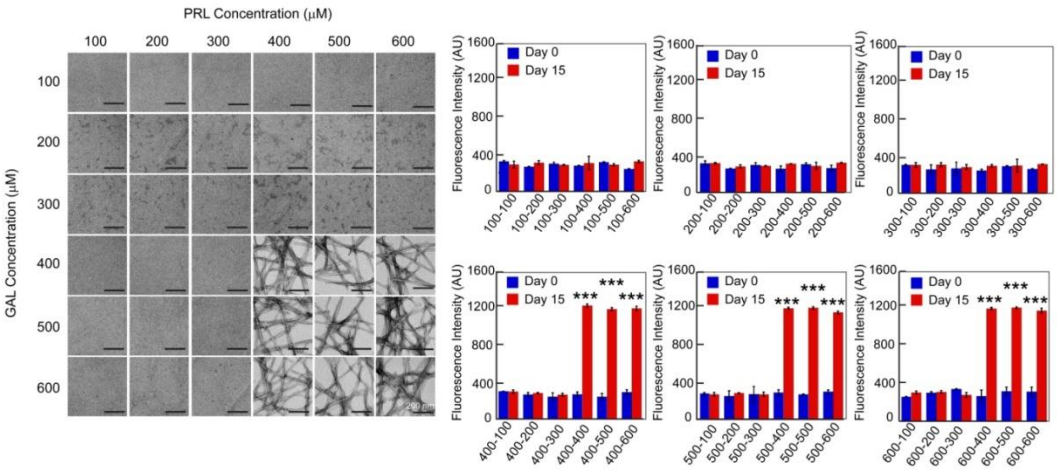


**Supplementary Fig 5: Concentration regime of PRL-GAL co-aggregation**. TEM images and ThT fluorescence of PRL-GAL co-aggregation at a varying concentration of PRL and GAL are shown. Both the (***Left panel***) TEM images and ThT fluorescence intensity at 480 nm (***Right panel***) suggest that PRL-GAL co-aggregation initiates above 400 µM concentration of either PRL/GAL as evident by the fibrillar morphology or high fluorescence intensity at 480 nm. Values represent mean ± SEM for n=3 independent experiments. The statistical significance(***p $\leq$ 0.001) is calculated by one-way ANOVA followed by the SNK post hoc test with a 95% confidence interval.

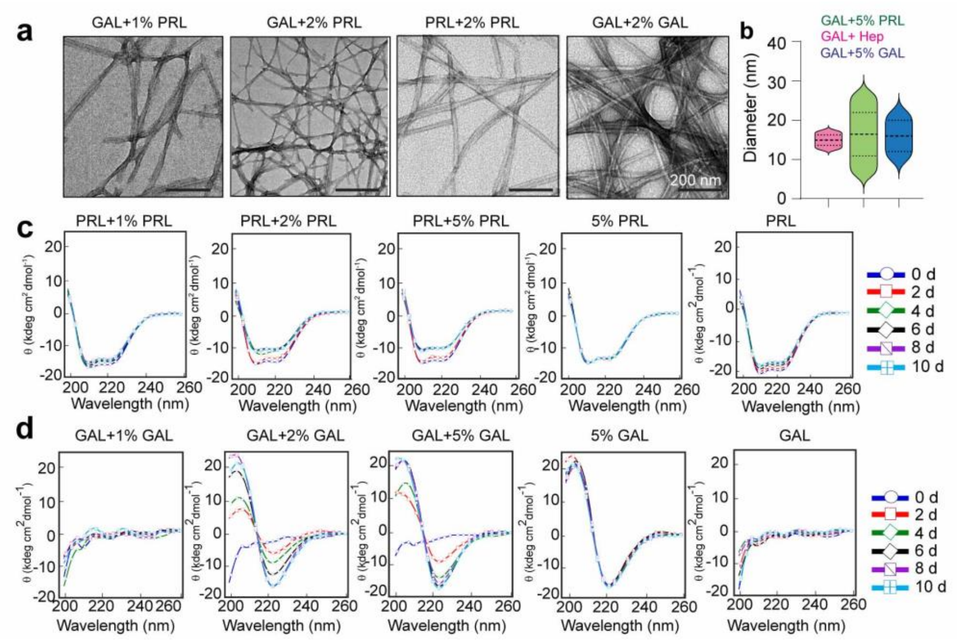


**Supplementary Fig 6: Aggregation and secondary structural transformations during seeding of PRL and GAL fibrils. a.** TEM imaging showing the presence of amyloid fibrils of GAL when seeded with 1% and 2% (v/v) PRL seeds, respectively and presence of amyloid fibrils of PRL and GAL when seeded with 2% (v/v) PRL and GAL seeds, respectively. **b.** Median values of different fibril diameters are shown with violin plots. **c.** CD spectra show the change in molar ellipticity of α-helical secondary structure of PRL in the presence of 2% and 5% PRL seeds whereas 1% PRL seeds show no change in secondary structure. 5% PRL seed and PRL alone retain their secondary structure after incubation. **d.** CD spectra show the change in GAL secondary structure from RC to β-sheet in presence of 2% and 5% GAL seeds whereas 1% GAL seeds show no change in secondary structure. GAL alone or only 5% GAL seed and GAL alone retain their secondary structure after incubation.

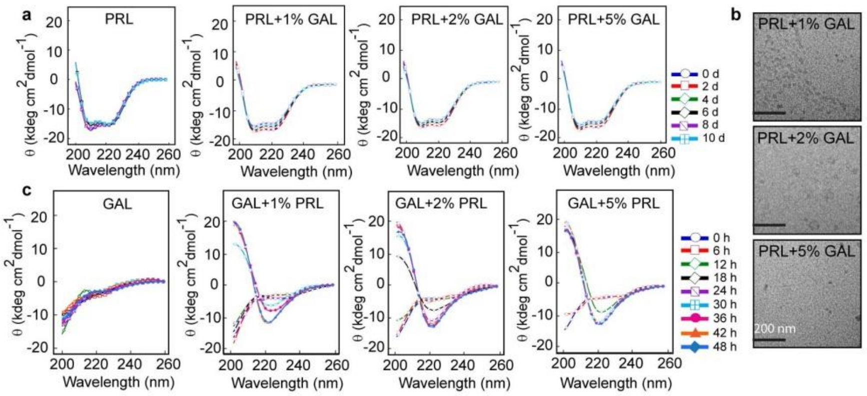


**Supplementary Fig 7: Secondary structural transformations due to cross-seeding of PRL and GAL fibrils. a.** CD spectra show no change in the α-helical secondary structure of PRL in the presence of a different concentration of GAL seeds (1%, 2%, and 5%). b. TEM images showing no fibril formation by PRL in the presence of 1%, 2%, and 5% GAL seeds. Scale bar is 200 nm as shown. **c.** CD spectra showing changes in the secondary structure of GAL from RC to β-sheet in presence of different PRL seed (1%, 2%, and 5%) concentrations**.**

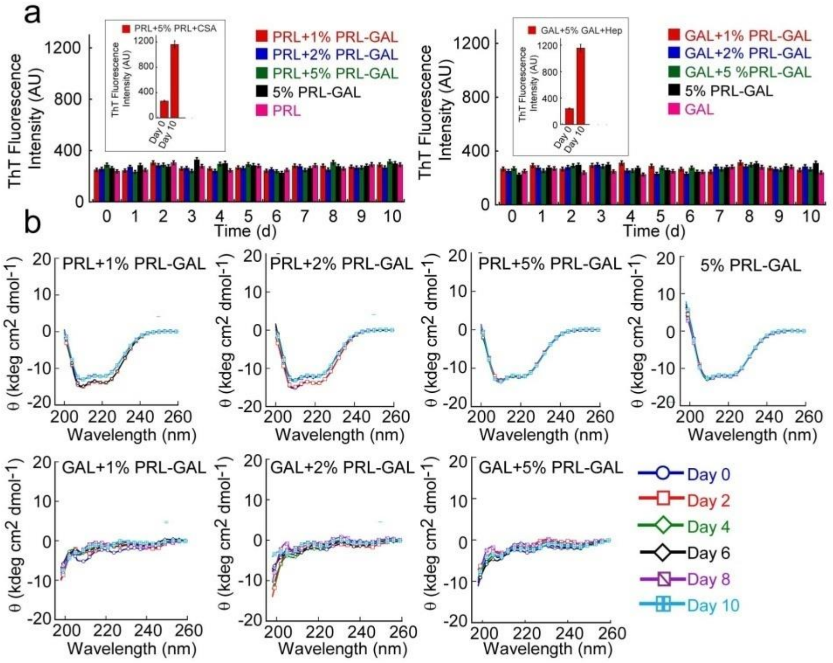


**Supplementary Fig 8: Cross-seeding of PRL and GAL by PRL-GAL co-fibrils. a.** ThT fluorescence intensity values with time indicating no aggregation of PRL or GAL in presence of different concentrations of PRL-GALfibrils seeds (1%, 2%, and 5% v/v). Only PRL-GAL seed and PRL/GAL monomer act as controls, which also showed no ThT binding. **b.** CD spectra showing no change in the α-helical secondary structure of PRL or random-coil structure of GAL in presence of a different concentration of PRL-GAL fibril seeds (1%, 2%, and 5%).

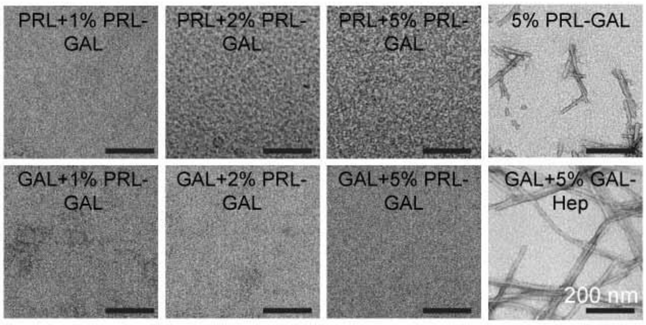


**Supplementary Figure 9**. **Electron microscopy of cross seeding of PRL and GAL by PRL-GAL co-fibril seed.** TEM images showing no fibril formation by either PRL or GAL in presence of 1%, 2%, and 5% PRL-GAL seeds. Scale bar is 200 nm as shown.

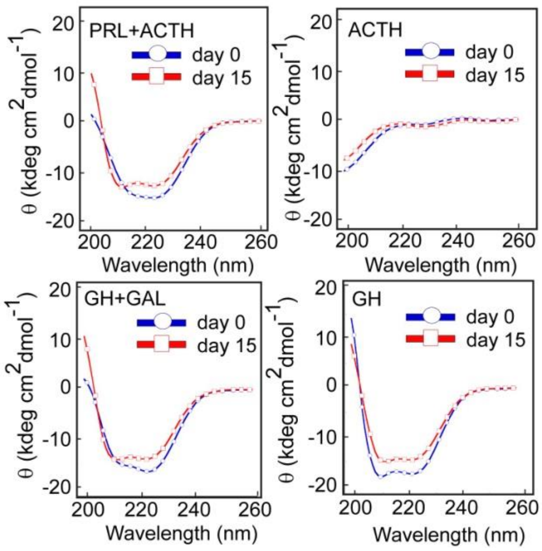


**Supplementary Fig 10: Conformational transition by different pairs of hormone co-aggregation using CD.** CD spectra showing no significant secondary structural transformation of PRL-ACTH and GAL-GH indicating no synergistic aggregation of these hormone pairs. The experiment is performed three times with similar results.

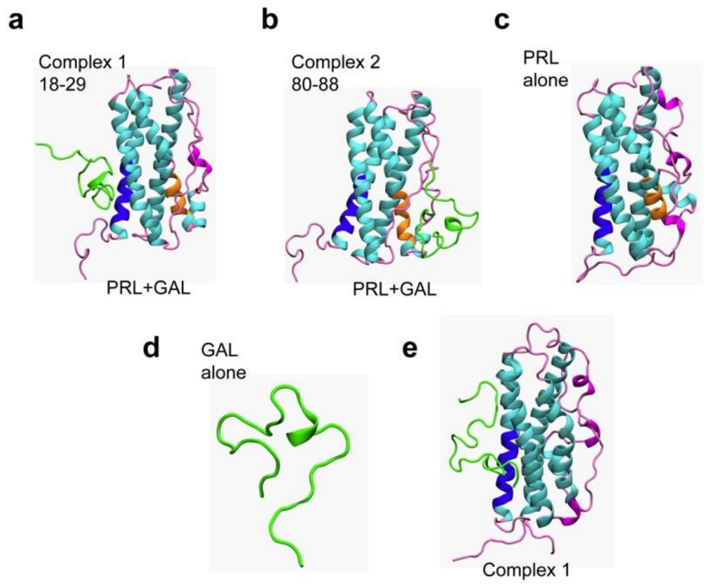


**Supplementary Fig 11: Protein-protein docking and MD simulations of PRL-GAL complexes. (a-b).** Structures of PRL-GAL docked complexes from protein-protein docking studies, which were subsequently used as the starting structures for MD simulations. The two lowest energy PRL-GAL complexes as obtained from docking studies are defined as follows – complex-1: GAL docked at PRL residues 18-28 (blue); complex-2: GAL docked at PRL residues 80-88 (orange). GAL is shown in green and PRL helices are shown in cyan and loops-coils are in magenta. **c.** Snapshots from MD simulations of individual PRL. **d.**Snapshots from MD simulations of individual GAL. **e.** The structure of complex-1 showing no change in either of the PRL/GAL structures after simulation using amber ff99SB force field.

Supplementary Table 1: Protein sequences used in the study

| Protein | Sequence |
|---|---|
| Human prolactin (PRL) | LPICPGGAARCQVTLRDLFDRAVVLSHYIHNLSSEMFSEFDKRYTHGRGFITKAINSCHT SSLATPEDKEQAQQMNQKDFLSLIVSILRSWNEPLYHLVTEVRGMQEAPEAILSKAVEIE EQTKRLLEGMELIVSQVHPETKENEIYPVWSGLPSLQMADEESRLSAYYNLLHCLRRDS HKIDNYLKLLKCRIIHNNNC |
| Human galanin (GAL) | GWTLNSAGYLLGPHAVGNHRSFSDKNGLTS |
| Human growth hormone (GH) | FPTIPLSRLFDNAMLRAHRLHQLAFDTYQEFEEAYIPKEQKYSFLQNPQTSLCFSESIPTP SNREETQQKSNLELLRISLLLIQSWLEPVQFLRSVFANSLVYGASDSNVYDLLKDLEEGIQ TLMGRLEDGSPRTGQIFKQTYSKFDTNSHNDDALLKNYGLLYCFRKDMDKVETFLRIVQ CRSVEGSCGF |
| Human adrenocorticotropic hormone (ACTH) | SYSMEHFRWGKPVGKKRRPVKVYPNGAEDESAEAFPLEF |
